# Unravelling the influence of light on inshore coral and sponge recruits and their substrate communities

**DOI:** 10.1101/2025.08.05.668793

**Authors:** Gerard F. Ricardo, Muhammad Azmi Abdul Wahab, Eduardo Arias, Lee Bastin, Christopher A. Brunner, Heidi M. Luter, Matthew Nitschke, Matt Salmon, Andrew P. Negri

## Abstract

Recruitment of progeny to coral reef populations involves complex ecological interactions, influenced by environmental factors such as altered underwater light conditions associated with poor water quality. Here, we exposed newly settled corals (*Acropora millepora* and *Acropora cf. tenuis*), the sponge (*Phyllospongia foliascens*), and their substrate communities to various light intensities and spectral profiles relevant to turbid inshore reefs. Coral and sponge recruit survivorship and growth generally exhibited an inverted U-shaped response to light intensity, suggesting environmental optima at lower light levels, while the influence of light spectra remained less clear within environmentally realistic treatment combinations. Crustose coralline algae cover similarly peaked at lower light levels, whereas turf algae increased with higher light conditions. Competitive interactions between the substrate communities and the recruits, along with photophysiological responses, were also assessed. Our results suggest that altered light characteristics associated with turbidity may not be as detrimental to coral and sponge recruits as other sediment-related stressors.

## Introduction

The capacity of coral reefs to recover following disturbances has become an area of increased research focus as intensified disturbances are predicted in the coming decades from a range of global and local impacts, including climate change and water quality degradation (Hughes et al. 2017a, Hughes et al. 2017b, Donovan et al. 2021). Recolonisation of the reef substrate under these altered conditions is critical for the future state of reefs, which may shift to alternate community assemblages following environmental stress events (McCook 1999, McManus & Polsenberg 2004, Johns et al. 2018, Hughes et al. 2019). Therefore, understanding the influence of pressures such as degraded water quality conditions, which often correlates with poor recruitment of colonisers, is vital (Thomson et al. 2021). Relative to their size, inshore reefs are disproportionately important economically, culturally, and recreationally, and often subject to the widest range of water quality conditions (Commonwealth of Australia 2023). On the Great Barrier Reef (GBR), inshore reefs are exposed to terrestrial sediment through riverine discharge, natural resuspension events and coastal development activities such as dredging (Fabricius 2005, Wolanski et al. 2005, McCook et al. 2015). Terrestrial runoff also contributes dissolved and particulate nutrients, particularly during periods of intense rainfall and flooding (Devlin & Brodie 2005, Kroon et al. 2012). Such exposures affect reef taxa through multiple mechanistic pathways, including suspended and deposited sediments, as well as alterations in both the quantity and quality of available light, which can lead to a range of physiological responses, including reduced photosynthesis, impaired gas exchange, smothering, tissue damage and necrosis, elevated energy expenditure, and increased susceptibility to bleaching (Jones et al. 2016, Tuttle & Donahue 2022). Early life stages are particularly susceptible to sediment exposures through the fertilisation, settlement and postsettlement phases (Babcock & Smith 2002, Fabricius et al. 2003, Ricardo et al. 2018, Brunner et al. 2021).

Among the mechanistic pathways associated with sediment and nutrient exposure, the least explored is how changes in the underwater light spectral profile caused by suspended particulates affect benthic reef taxa. A range of interacting biotic and abiotic factors including river discharge, wind-driven resuspension, coastal development, plankton dynamics, and flocculation processes influence the concentration and composition of suspended particles in the water column, leading to shifts in the light spectral profiles experienced by autotrophic reef taxa (Storlazzi et al. 2015, Maggiorano et al. 2025). In clear shallow-water environments, longer wavelengths of the visible electromagnetic spectrum (reds and oranges) are attenuated, with only shorter wavelengths remaining, primarily greens and blues, as depth increases (Dustan 1982). Additionally, increases in particulates such as sediment grains and planktonic algae scatter, absorb and attenuate light causing shifts in the spectral profile to the centre of the visible spectrum resulting in yellow-green downwelling irradiance (Strydom et al. 2017b). Many benthic taxa rely on symbiotic relationships with photosynthetic microorganisms, which provide a significant portion of the host’s energy requirements through autotrophic carbon fixation (Wilkinson 1983, Muscatine 1990). However, benthic autotrophs primarily utilise specific segments of the visible spectrum (known as Photosynthetically Usable Radiation (PUR)), owing to the specificity of their pigments. For example, chlorophyll *a* pigments generally absorb blue light (∼ 440 nm) and red light (∼660 nm) more efficiently than mid-range (yellow-green) wavelengths associated with waters affected by suspended sediments and planktonic algae (Morel 1978, Dubinsky et al. 1984).

In addition to changes in the light spectral profile, changes in water quality attenuate light through absorption and scattering. For example, in the absence of water column particles, light at the water’s surface is attenuated by ∼66% at 5 m depth, and to ∼86% at 10 m depth (Jones et al. 2020). However, depth-attenuation effects are further compounded by particles via scattering and absorption in turbid environments causing coastal darkening, with the photic zone of nearshore reefs reduced to shallower areas (Magno-Canto et al. 2019, Morgan et al. 2020, Wernberg & Straub 2024). Previous studies have identified key light thresholds for zooxanthellate corals, with a 5.2 mol photons m^-2^ d^-1^ (Muir et al. 2015), and 7 to 8 mol photons m^-2^ d^-1^ estimated as the minimum required to sustain reef accretion (Kleypas 1997). Experimental research indicates that coral light limitation thresholds vary considerably, ranging from 2 to 16 mol photons m^-2^ d^-1^ (reviewed by Canto et al. (2021)).

Coral reef substrate colonisers, such as corals and sponges, play key ecological roles in reef environments, contributing to structural complexity, habitat provision, nutrient cycling, and species interactions through both competition and facilitation (Bell 2008, Brandl et al. 2019). Early benthic life stages of these taxa are particularly vulnerable to changes in light quantity and quality, as these stages often rely on symbiont-derived autotrophy for energy. Many corals begin the benthic stage of their lifecycle without, or with few, algal symbionts i.e., horizontal transmission (Figueiredo et al. 2012, Toh et al. 2013), and typically settle in crevices that are limited in light (Nozawa et al. 2011, Ricardo et al. 2021). However, following settlement, photosynthetic symbionts of the family Symbiodiniaceae infect and increase in density, leading to a shift towards autotrophy that can eventually be responsible for >80% of the colony energetic needs (Muscatine 1990, Brunner et al. 2022). For example, photosynthetic symbiont density increases in newly settled *Acropora* recruits rapidly between 40 and 71 days (Quigley et al. 2019), but the extent to which recruits rely on existing energy reserves, feed heterotrophically, or shift to autotrophy as the dominant energy pathway is poorly understood and likely varies between season, species and reef position along the shelf i.e., inshore vs offshore (Anthony 2000, Toh et al. 2013, Randall et al. 2020). Importantly, the timing of this shift to autotrophy will determine whether early-stage recruits are affected by changes in light during their initial developmental period.

Similar to corals, photosynthetic symbionts in reef sponges can provide substantial nutritional requirements of the sponge host (Wilkinson 1983). Sponge larvae acquire symbionts primarily via vertical transmission, though some species also incorporate symbionts from the environment (horizontal transmission) (Maldonado 2007, Schmitt et al. 2008). Notably, the common Indo-Pacific sponge *Phyllospongia foliascens*, which inhabits but is not exclusive to inshore reefs, is a gonochoric species harbouring cyanobacterial symbionts (e.g. *Synechococcus* spp.) that are vertically transmitted from the maternal parent to the offspring (Usher et al. 2005, Luter et al. 2020). This early establishment of photosynthetic symbiosis may increase its susceptibility to low or altered light conditions during recruitment. Relatively little is known about tropical sponge postsettlement processes and research in this area lags behind other well-studied taxa such as corals.

The recruitment of corals and sponges involves a complex relationship with the encrusting and turf algae that colonise the coral reef substrate. Among the most ecologically important encrusting taxa are crustose coralline algae (CCA), which are crucial framework builders and play a pivotal role in coral larval settlement in addition to other invertebrates (Rodriguez et al. 1993, Harrington et al. 2004). CCA can also act as a facilitator or competitor during postsettlement survival of recruits (Brunner et al. 2022). One of the most dominant genera on the GBR is *Porolithon*, common in upper shelf and reef flats (Harrington 2004, Dean et al. 2015), and an effective settlement inducer for corals of the genus *Acropora* (Harrington et al. 2004, Whitman et al. 2020). Established crusts of *Porolithon spp.* are sensitive to low-light periods (Bessell-Browne et al. 2017); however, early-stage germlings and growth may have different susceptibilities to changes in light-environment than more established CCA (Ordoñez et al. 2017). CCA are also sensitive to deposited sediments (Harrington et al. 2005, Ricardo et al. 2021) and CCA abundance is negatively correlated with turbidity across inshore-offshore gradients (Fabricius & De’Ath 2001). In contrast, filamentous algal turfs (often defining a broad group of mostly filamentous macroscopic algae <2 cm, such as benthic diatoms, cyanobacteria, *Chlorophyta*, *Rhodophyta* and *Phaeophyta*; Connell et al. 2014; Tebbett & Bellwood 2019) are reported to have negative associations with recruiting sessile invertebrates (Birrell et al. 2005, Cárdenas et al. 2016). These algae rapidly colonise substrates after disturbances and often thrive in inshore-reef conditions such as increased nutrients, sediments, and alternate light regimes (McCook et al. 2001, Birrell et al. 2005, Tebbett & Bellwood 2019).

To better understand the relationship between these early-stage substrate colonisers and light quality and quantity, we first characterised the light environment of an inshore reef site based on *in situ* measurements. We then established irradiance–response relationships in accord with the ANZECC water quality guidelines (Warne et al. 2018), across a range of Daily Light Integrals (0 to 9 DLI) under broad-spectrum light profiles typical of clearer inshore tropical waters, and examined how these relationships change under shifted-spectrum light profiles representative of turbid conditions. Common aquarium lighting used in experimental light studies often exploit PUR by providing spectra rich in the blue and red ranges, thereby reducing energy costs associated with emitting less photosynthetically efficient wavelengths. However, such lighting may not accurately represent natural light environments, particularly for species capable of pigment plasticity or spectral acclimation. Here we use a novel spectrally tuneable lighting system to experimentally manipulate the underwater light environment with high precision, allowing us to simulate realistic spectral conditions observed on turbid inshore reefs and assess their effects on early life stages of benthic autotrophs. As shifts in the light spectral profile co-occur with light attenuation *in situ*, we used a crossed experimental design to decouple these variables.

## Methods

### *In situ* measurements for experimental treatments

To first determine realistic light spectral profiles of inshore reefs, *in situ* light measurements were collected over a two-month period in 2018 at an inshore site (Florence Bay; 19.121°S, 146.882°E) within Cleveland Bay, GBR, Australia. Measurements were taken using an eight-wavelength (425, 455, 485, 515, 555, 615, 660, 695 nm) multispectral radiometer (MS8, In-situ Marine Optics, Australia) positioned adjacent to a nephelometer (DL3, In-situ Marine Optics). Median depth was 4.6 m during the deployment (2.4 m relative to the lowest astronomical tide). After retrieval of the sensor, the irradiance data in watts (µW cm^-2^ nm^-1^) were converted to photons (µmol photons m^-2^ s^-1^ nm^-1^) using the equation (Thimijan & Heins 1983, Ricardo et al. 2021):

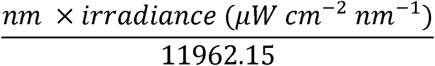

To calculate PAR, a spline was fit to individual measurements and the area under the curve between 400 and 700 nm calculated using the package *MESS* (Ekstrom 2014) in *R* (v. 4.0.2; R Core Team (2013)). To calculate DLIs, the daily PAR values were fit using a loess spline with the package *stats*, predicted for each second, and summed. A turbidity index (blue:green ratio) was calculated following Jones et al. (2021), using the ratio of irradiance at 455 nm (i.e., blue) to that of 555 nm (i.e., green). As green wavelengths are more characteristic of turbidity, higher values indicate clearer water whereas low values indicated turbid water. Values derived from the water-quality light data were used to inform the experimental design explained below.

### Species collection, larval culturing, and substrate conditioning

The coral genus *Acropora* is the main framework builder and coloniser after disturbance on the GBR and the Indo-Pacific (Sheppard et al. 2008, Ortiz et al. 2021), and two species, *Acropora millepora* (Ehrenberg, 1834) and *Acropora cf. tenuis* (Dana, 1846), were selected to represent this genus. Coral collection, spawning, larval rearing and substrate conditioning followed Ricardo et al. (2021) and Heyward and Negri (1999). Briefly, gravid adult colonies of each species were collected at 5–8 m depth from Backnumbers Reef (18°30’47”S, 147°09’08”E; mid-shelf GBR) and transported to the National Sea Simulator at the Australian Institute of Marine Science (AIMS), Townsville in November 2018. On the nights of spawning, egg-sperm bundles from six to eight colonies were collected and cross-fertilised. The embryos were then washed free of sperm twice and transferred into 500 L indoor flow-through fiberglass tanks to undergo embryogenesis and larval development.

Artificial substrates for coral settlement and recruit grow-out were comprised of 7-cm diameter polyvinyl chloride (PVC) discs. PVC was selected as a substrate material as it is highly successful for colonisation of biological settlement inducers (Kennedy et al. 2017, Ricardo et al. 2017), and can easily be manufactured to prevent light-confounding effects i.e., shading. Each disc was conditioned in flow-through outdoor aquaria for ∼3 months under 70% block shade cloth, which reduced maximum daily light intensity to ∼250 µmol photons m^-2^ s^-1^. *Porolithon* spp., a common crustose coralline algal genus known to induce settlement of *Acropora* coral larvae (Heyward & Negri 1999, Whitman et al. 2020), was added to the conditioning tanks. Conditioning of CCA onto the substrates prior to experiment was ∼10% of the substrate area. A total of 597 *A. millepora* recruits and 480 *A.* cf. *tenuis* recruits were used in the experiment.

Thirty gravid female *P. foliascens* were collected from depths of <3m from an inner-shelf reef (Fantome Island; 18°41’10”S, 146°30’54”E), in early November 2018, and transported to the National Sea Simulator, AIMS. Environmental and tank conditions for adult sponge husbandry, as well as specific methodology for larval collection are described in detail in Abdul Wahab et al. (2019). Larvae used for settlement and the production of sponge recruits in this study were pooled from at least 10 female sponges. Settlement substrates comprised of white aragonite plugs (2 cm diameter), which have been found suitable for sponge settlement, and were conditioned for two weeks in a similar light conditions used for the adult sponges but receiving flow-through unfiltered seawater.

As *P. foliascens* larvae are negatively phototactic (Abdul Wahab et al. 2019), settlement was directed onto shaded, upward-facing surfaces by positioning preconditioned aragonite plugs upside-down under overhead lighting that mimicked typical adult light conditions. Larvae (24 h old; n = 50–70) were added to each container and incubated for 24 h, after which settled and metamorphosed recruits were transferred to flow-through unfiltered seawater for a two-week acclimation period prior to the experiment. A total of 945 *P. foliascens* recruits (2–3 weeks of age post settlement) were available after the two-week acclimatisation period.

### Experimental design and set-up

Spectrally tuneable LED light fittings (520 × 430 mm) were designed in-house to simulate realistic light spectral profiles found in marine environments (described in Luter et al. 2021). Briefly, each light contained 840 individual LEDs controlled by 19 independent colour channels. The lighting system was integrated into a programmable-logic-controller (Siemens PCS7, Supervisory Control and Data Acquisition System), with the channels adjusted through a feedback system to a user-defined light-intensity level. Neutral-density shade film was also added to the lower-light treatments to allow for greater control of the colour channels. A PAR Quantum sensor (Skye SKP 215) was positioned on the grid floor of the tank and connected to the programmable logic controller (PLC) for feedback, allowing the PLC to control the light intensity. While the lights were designed to provide wavelengths in the visible spectrum (400–700 nm), a small amount of light in the UV-A range (370–400 nm) was also present.

The light experiment was conducted under nine LED lighting systems mounted above large temperature-controlled water baths (1,200 L volume; *n* = 9). Each water bath housed two replicate 50 L experimental tanks (*n* = 2 per bath), which were placed half-submerged to maintain thermal stability. This resulted in a total of 18 experimental tanks. The experimental design included five light intensity levels applied across two spectral profiles (broad and shifted), with details provided below (Fig. S1 a,b). However, because there is no spectral difference at complete darkness (0 DLI), only one shared treatment was used for that condition, resulting in nine lighting/water bath systems in total rather than ten. All recruit substrates were positioned so they were within ∼10% of the maximum light intensity (Fig. S1 c).

General purpose and raw seawater supplied to the experimental room were temperature controlled to within 0.3 °C of the designated setpoint of 27 °C. While the experimental tanks were not connected to an automated temperature control system, the use of the large water baths and room air-conditioning (set to 28 °C, allowing for 1 °C of evaporative cooling) helped buffer temperature fluctuations. All tanks were manually checked with a temperature probe prior to commencement of the experiment, and three randomly selected water baths were logged for several days to ensure no temporal temperature deviations (mean ± SD = 27.15 ± 0.26 °C). A storm-related issue with the raw-water feed into the broad-spectrum 1 DLI tanks led to a clear, isolated effect. As the first outlet in the circular seawater outlet system, it received water with visible turbidity following the storm, while the remaining tanks remained unaffected. Subsequently this treatment, which included both corals and sponges, was removed from all analyses.

Based on *in situ* field data, the experiment was designed to examine the effects of low light intensity (5 levels) and spectral quality (broad and shifted spectrums). Coral recruits tend to settle in crevices and grooves, and therefore growth *in situ* over the first 6-weeks following settlement are likely to be in low-light environments (Doropoulos et al. 2016, Ricardo et al. 2021). Therefore, we conservatively consider the expected range of light intensities for early coral recruitment to be 0 to 100 µmol photons m^-2^ s^-1^ (0 – 2.5 Daily Light Integrals (DLI)). Less is known about typical light levels for sponge recruits, but they are assumed to fall within a similar range. Five light intensities were selected to cover the range of environmentally realistic values. Light intensities followed a sinusoidal ramping over the course of the day (06:00 to 18:00) with peak midday PAR levels approximately 0, 10, 30, 100, 300 µmol photons m^-2^ s^-1^, which equated to DLIs of approximately 0, 0.3, 0.9, 3, 9 mol photons m^-2^ d^-1^. For corals, these values relate to PUR DLIs of 0, 0.14, 0.51, 1.65, 5.06 mol photons m^-2^ d^-1^ for the broad spectral profiles, and 0, 0.04, 0.38, 1.32, 3.81 mol photons m^-2^ d^-1^ for the shifted spectral profiles based on the absorption coefficients of *Symbiodinium* in Hennige et al. (2009).

Using the blue:green ratios described above, two treatment levels were selected based on *in situ* measurement ratios (i.e., Turbidity Index; Fig. 1a), which equated to ∼0.8 and ∼9 NTU (Nephelometric Turbidity Units). The broad-spectral profile was slightly right-skewed and had a dominance of wavelengths in the 400 to 600 nm range (violet, blue and green), whereas the shifted spectral profile was normally distributed and had a dominance of wavelengths in the 500 to 600 nm (green–yellow) range, and visually the light appeared yellow (Fig. 1 e,f). To determine causative effects, we employed a crossed design which resulted in both realistic and unlikely environmentally relevant conditions. Since light intensity and blue:green ratios are typically correlated *in situ*, some pairings such as high light with a low blue:green ratio, are uncommon. Specifically, the shifted-spectral profile we used to simulate chronic turbidity (∼9.4 NTU; blue:green ratio of ∼0.23) is considered relatively high in *in situ* field conditions, with the blue:green ratio constrained by light intensity occurring only up to DLIs of 3.4 mol photons m^-2^ d^-1^ (99^th^ percentile, Fig. 1 a,b).

**Fig. 1.**
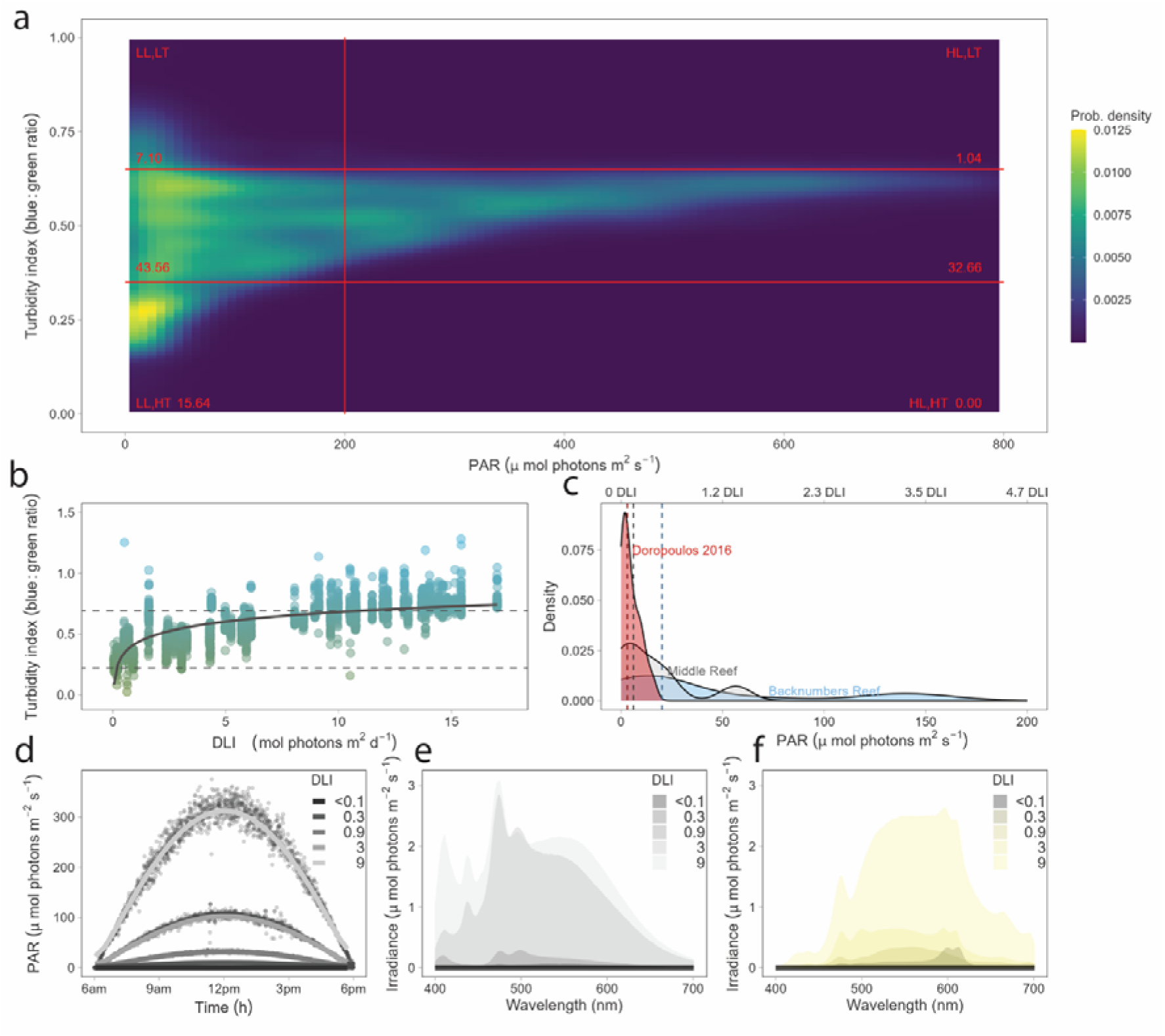
*In situ* light environment of a lower-turbidity inshore reef, and experimental laboratory conditions used during the six-week light exposure on coral recruits. (a) Probability densities of light intensity and the turbidity index (blue (455 nm) to green (555 nm) ratios) during daylight hours at Florence Bay in 2018 over ∼65 d. Lines between 0.35 and 0.65 blue:green ratio, and 200 µmol photons m^-2^ s^-1^ (red lines) were used to summarise probabilities (%) within these regions (red text). Low and high light intensities and turbidity denoted by LL, HL, LT, HT in the plot. (b) The relationship between daily light integrals (DLI) and turbidity index for this deployment. Dashed lines indicate blue:green ratios used in the experimental treatments. (c) Light intensity levels (Photosynthetic Active Radiation; PAR), suitable for recruitment, were measured at midday in grooves at both an inshore and offshore reef site, compared with data from Doropoulos et al. (2016). Experimental light exposures of a representative day: (d) PAR and DLIs with points representing PAR measurements and fitted lines used to derive DLIs, (e) relative broad-band spectral profiles representing ∼0.8 NTU, and (f) relative shifted spectral profiles representing ∼9.4 NTU. Each spectral profile was binned to 1 nm wavelength bands.

Irradiance in the experiment were measured with a Jaz spectrometer (Ocean Optics) calibrated to a recently calibrated HL-2000 lamp. The spectrometer was connected to a 5 m long 400-µm diameter fibre optic with a 3.9 mm diameter in-water cosine corrector attached. Irradiance from spectrometers was interpreted as relative rather than absolute values because of the common issues associated with spectrometer calibrations, light flickering, and attenuation of light under cosine correctors (Johnsen 2012), and were therefore corrected against the PAR Quantum sensor. All absolute irradiance readings were converted to PAR using the described equation above.

### Coral recruit light-exposure experiment

Newly settled corals were introduced to experimental treatments at the start of the day while light levels were low and reflected their likely settlement *in situ* light environment (< 50 µmol photons m^-2^ s^-1^) (Nozawa et al. 2011). Three discs of each species were placed in each of the two experimental tanks. For *A. millepora* (median = 7; range: 1–25) and *A.* cf. *tenuis* (median = 5; range: 1–44), this resulted in ∼50 recruits per treatment combination. The recruits were infected with cultured Symbiodiniaceae species *Cladocopium proliferum* (AIMS ID: SCF055-01.10) (Butler et al. 2023), in a designated symbiont infection tank (1–2 × 10^4^ cells L^-1^) on days 6 and 12 after settlement. These symbionts were cultured under the conditions: 12:12 cycle; 60 μmol photons m^−2^ s^−1^, 27 °C).

A small proportion of raw (unfiltered) seawater was mixed with FSW for heterotrophic feeding (Conlan et al. 2017), and the total mix pumped into each flow-through tank at ∼0.75 L min^-1^. The substrates were transferred to new 50 L tanks every 4 days to reduce excessive algal growth, and each substrate was also lightly wiped with a soft paintbrush to control algae growth. While these measures were used to manage algae growth, they did not entirely remove it, as algae-coral competition was an important response under various light regimes. The substrates were photographed at settlement and following the six-week exposure with high resolution images using a Nikon D810 with a Nikon AF-S 60mm f/2.8G ED macro lens and four Ikelite DS161 strobes to determine survivorship and growth. Where two or more individuals had settled in a cluster, only the group was counted. Horizontal growth was measured using inbuilt tools in Photoshop (v. 20.0.4) on haphazardly selected individual recruits. Where partial mortality occurred, defined here as visible tissue loss, only the living section of the recruit was counted. While horizontal growth is a well-established method for assessing growth in coral recruits (Bak & Engel 1979, Vermeij 2006, Doropoulos et al. 2012), vertical growth from the initial polyp also occurs in the first weeks but is difficult to measure. As a metric of early development, the estimates of growth made from horizontal growth are likely to be conservative.

Changes in the photophysiology of the corals were measured with Pulse Amplitude Modulating (PAM) fluorometry (corals: IMAGING-PAM MAXI, Walz; CCA: MINI-PAM, Walz) using dark-adapted quantum yields (*F_v_/F_m_*) and rapid light curves. The minimum and maximum dark acclimated fluorescence yields, given by *F_0_* (minimum fluorescence) and *F_m_* (maximum fluorescence) respectively, are used to calculate *F_v_* (variable fluorescence), where *F_v_* = *F_m_* – *F_0_*. For *F_v_/F_m_*, corals were left in darkness following a night-time dark cycle with sampling on haphazardly selected substrates commencing at 08:00. The minimum fluorescence signal was kept above 130 units to ensure an adequate signal-to-noise ratio (Ralph et al. 2015). Rapid light curves describe the photosynthetic capacity of the organism, its light adaptation state, and its capacity to tolerate short-term changes in light whereas dark-induction curves provide information on changes in quenching responses over time (Ralph & Gademann 2005). For all rapid light curves, measurements were collected bracketing midday (all lights held constant during analysis and substrates selected haphazardly) and dark-adapted for 2 minutes. Ralph and Gademann (2005) recommends a 5–10 s quasi-darkness to allow the rapid oxidation of the primary electron acceptor (*Q_A_*) but this short period was not possible. A stepwise series of ten light intensities (450 nm, min = 0 μmol photons m^−2^ s^−1^; max = 930 μmol photons m^−2^ s^−1^) were applied progressively increasing every 10 s to determine the relative electron transport rate (*rETR*) calculated using (PAR × Δ*F*/*F_m_*’), and all *rETR* data were fit to a standard double exponential decay function Platt equation (Platt et al. 1981, Ralph & Gademann 2005) in the software R (v. 4.0.2). The parameters α (initial slope, photosynthetic efficiency), *E_k_* (the minimum saturating irradiance; photochemical quenching dominates below *E_k_*, while nonphotochemical quenching dominates fluorescence quenching above *E_k_*), *rETR_max_*(the maximum relative electron transport rate) and *E_m_* (light intensities that correspond to the *rETR_max_*) were derived from the model (Ralph & Gademann 2005) (Fig. S2a). Poor model convergence of the Platt equations and high mortality on certain substrates resulted in missing data for some treatment levels. However, this is unlikely to have substantially affected the observed trends, as the remaining treatment levels still spanned the most environmentally realistic conditions.

The alternate spectral profiles, in addition to corallite skeletal structure, can influence acclimation states and light absorption. As such, we additionally derived *E_k_* from:

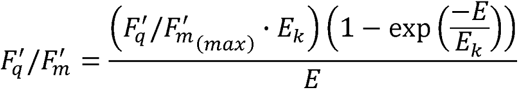

as a second validation of the acclimation state per spectra (Hennige et al. 2008, Nitschke et al. 2018). *E_k_* derived from this equation (light-dependent quantum efficiency of PSII) correlated well with that of the Platt equation (r^2^ = 0.733) (Fig. S3).

To additionally assess the poise of the photosystem, we calculated the extent of light-dependent photochemical quenching [1 − C] and non-photochemical quenching [1 − Q] from each rapid light curve step as follows:

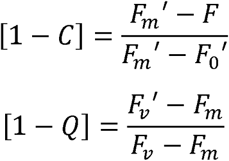

Here, [1 − C] is equivalent to *qP* in other studies and assumes zero connectivity between adjacent reaction centres and represents the proportion of excitation energy used in photochemistry, indicating the fraction of open reaction centres. [1−Q] quantifies dynamic non-photochemical quenching, which is equivalent to the excitation pressure over photosystem II (PSII) (Suggett et al. 2015, Nitschke et al. 2018). The product of [1 − C] and [1 − Q] accounts for total energetic dissipation (Nitschke et al. 2018).

### Sponge recruit light-exposure experiment

Immediately prior to introducing the *P. foliascens* recruits to the light treatments, substrates were distributed across nine separate racks corresponding to the experimental levels. Each rack held 21–24 individual substrates, totalling approximately 118 recruits per treatment combination. Sponge recruits were unevenly distributed among substrates (median = 4, range: 1–19). Each rack was placed in a 50 L experimental tank containing the coral substrates corresponding to the assigned light treatment. The survival of individual sponge recruits was recorded at Day 0, 3, 6, 10, 14, 21, 36 and 42 of the experiment, through direct observations using a stereo microscope. The majority of algal growth was gently removed from each substrate using fine forceps and a pipette to control for overgrowth at each sampling time point. High resolution images of the substrates were captured using a Canon 7D Mark II with an EF 100 mm f/2.8 macro lens at Day 0 and 42 to quantify horizontal growth of the sponge recruits. Similar to corals, horizontal growth does not capture changes in vertical growth that may occur initially in sponge recruits, and therefore estimates of body size increase are likely to be conservative.

### Substrate community analyses

The substrate community was photographed with the coral and sponge recruits, and segmented using Trainable Weka Segmentation in ImageJ (Arganda-Carreras et al. 2017) on a Dell R820 256 GB RAM High-performance Computer System (Fig. S4). This machine learning approach used the default classifier Fast Random Forest with 200 trees. For each substrate type, a range of substrates across each treatment (n = 10) were used to train the model into ‘CCA’, ‘turf algae’, ‘biofilm’, ‘bare space’, and ‘bleaching/glare’. ‘Bleaching/glare’ described areas with little distinctive characteristics to reliably determine the substrate, and these regions were not used in the analysis. Further, plugs that had greater than 20% of this class were not analysed. While visually different, ‘biofilm’ was pooled with ‘bare space’, as ‘bare space’ likely still contained some level of biofilm. The resulting model was then further retrained iteratively to remove false positives and false negatives before running on the remaining images. Changes in the photophysiology of CCA were evaluated as described above for coral recruits by haphazardly selecting an area of *Porolithon spp.* for PAM fluorometry. The measuring light used peaked at 660 nm (red), which is appropriate for pigments of CCA (Burdett et al. 2014).

### Statistical analysis

To derive the DLIs, instantaneous measurements of the PAR recorded at 10s intervals were fit with LOESS splines and the area under the splines were determined using the package *MESS* (see Fig. 1d) in the statistical software program *R* (version 4.0.3). To allow for inverted U-shaped trends and to enable estimation of effect concentrations, data were fit with generalized binomial nonlinear hormesis models under a Bayesian framework in the package *bayesnec* (Fisher et al. 2024), using the equation *ecxhormebc5*:

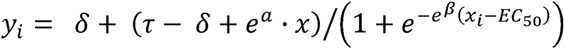

Model parameters included the upper and lower asymptotes (τ and δ), the slope (β), the effective concentration causing a 50% response (EC), and an adjustment term (α) allowing for asymmetry in the dose–response curve. Models were fit using 4 chains, with 5000 iterations (2500 warm up, 2500 sampling), and further iterations were added if required. A binomial error function was applied for binomial data (i.e., survivorship data), whereas a gamma distribution applied for continuous data with a zero asymptote (i.e., growth, rapid light curves etc.), and a beta distribution applied for proportion data (i.e., community composition). Each model was assessed using trace plots, and posterior predictive checks, all of which were satisfactory. Effect concentrations causing a 10% reduction (EC_10_), which are common effect-size threshold metric used in ecotoxicology, were calculated when possible (Warne et al. 2018).

#### Structural Equation Modelling (SEM)

To evaluate the underlying and indirect relationships between light spectrum, DLIs, the survival of coral recruits, and the cover of crustose coralline algae and turf algae, we employed a structural equation modelling (SEM) approach using the R package *lavaan*. The survival probability of coral recruits, proportion cover of CCA and turf algae were modelled as endogenous variables with their outcomes influenced by other variables in the system. Exogenous variables of DLI and light spectrum were included as predictors in the model, with DLI modelled as a continuous log_10_-transformed variable, and spectrum as a categorical variable (broad-spectrum used as the reference). Survival probability of coral recruits was modelled as a function of DLI, light spectrum, and their interaction. To capture non-linear effects, a quadratic term for DLI was included, along with interaction terms between DLI and spectrum. For the proportion of CCA and turf algae, the influence of DLI, light spectrum, and their interaction was tested. Survivorship observations were weighted based on their initial sample units. Additionally, we assessed the indirect effects of crustose coralline and turf algae cover on coral survivorship.

## Results

### *In situ* measurements for experimental treatment selection

The 65-day instrument deployment period was influenced by two large wind-driven natural resuspension events resulting in varied spectral profiles and a median DLI of 6.1 mol photons m^-2^ d^-1^. A DLI of 9 mol photons m^-2^ d^-1^ (used as the highest experimental light treatment) represented the 57^th^ percentile. The median PAR during daylight hours was 104 µmol photons m^-2^ s^-1^, and the median blue:green (455/555 nm) ratio was 0.53. The upper vertical tail of the probability density i.e., indicative of clearer water (Fig. 1a), was represented by blue:green ratios ≥0.65 and light levels ≤200 µmol photons m^-2^ s^-1^, which occurred for 7.6% of the daylight hours; whereas the lower vertical tail of the probability density i.e., indicative of turbid water was represented by blue:green ratios ≤0.35 and light levels ≤200 µmol photons m^-2^ s^-1^, which occurred for 15.7% of the daylight hours. Underwater light spectra corresponding to different turbidity levels and times of day are accessible at: ‘https://ricardo-gf.shinyapps.io/underwater_light_app’.

Consequently, the mean blue:green ratio used for the broad-spectral profile in the experiment was 0.72, whereas the mean blue:green ratio used for the shifted-spectral profile was 0.23 representing the lowest and highest levels of turbidity in Cleveland Bay. Comparison between the blue:green turbidity index and DLI revealed a nonlinear relationship, where DLI increased substantially with higher ratios. A 10% reduction in the blue:green ratio from the mean high DLI conditions occurred at 8.80 DLI, with this threshold indicating deteriorating water quality conditions (Fig. 1b). Light levels at settlement sites relevant to the recruits were typically very low at 20.5 µmol photons m^-2^ s^-1^ PAR for Backnumbers Reef and 6.0 µmol photons m^-2^ s^-1^ PAR for Middle Reef. These were in range of those measured at reef sites reported by Doropoulos et al. (2016) at ∼3 µmol photons m^-2^ s^-1^ PAR (Fig. 1c). As such, we considered the lower light treatments the most relevant to recruits within the initial postsettlement phase.

### Survivorship and growth

Across the range of exposures, the hormesis model resulted in an inverted U-shaped trend in *A. millepora* coral recruit survival in the broad-spectrum light profile, peaking at maximum = 0.18 mol photons m^-2^ d^-1^ (lower EC_10_ = 0.05, upper EC_10_ = 0.32, 95% CI: 0–0.53). There was also a stronger inverted U-shaped response under the shifted-spectral light profile, especially at higher light intensities (maximum survival = 0.17 mol photons m^-2^ d^-1^, lower EC_10_ = 0.04, upper EC_10_ = 0.45, 95% CI: 0–0.70) (Fig. 2a). Growth peaked between 0.3 and 1 mol photons m^-2^ d^-1^ (Fig. 2b) under the broad-spectrum light but was not affected by different intensities of shifted-spectrum light. The greatest growth occurred at 0.3 mol photons m^-2^ d^-1^ in the broad-spectrum with recruits 3-fold greater than their initial size.

**Fig. 2.**
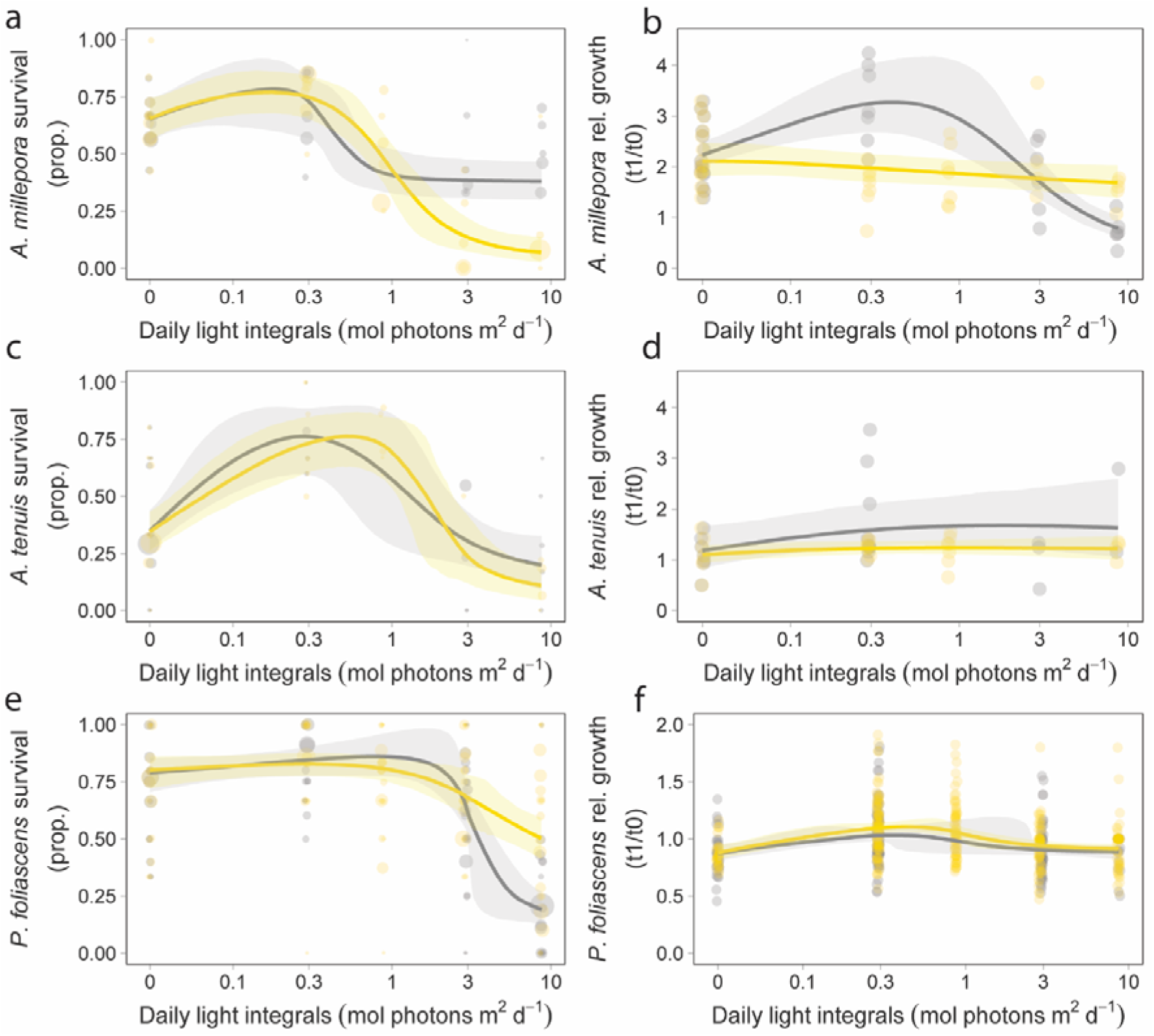
The effects of six-week light exposure under a range of light intensities (DLIs) and spectral profiles (broad and shifted) on the survivorship (a,c,e) and relative growth (b,d,f) of coral and sponge recruits. Survivorship and growth after the light exposure for (a, b) *Acropora millepora* coral recruits (c, d) *Acropora* cf. *tenuis* coral recruits, and (e, f) *Phyllospongia foliascens* sponge recruits. Grey line = broad-spectrum mean line; yellow line = shifted-spectrum mean line.

For the SEM analyses of *A. millepora*, the model structure aligned well with the observed data (p = 0.968), indicating a good overall fit (Fig. 3). Recruit survival was significantly influenced by both the linear (p < 0.001) and quadratic (p = 0.003) effects of DLI, indicating a unimodal response across the light gradient, consistent with the hormesis curves. A significant interaction between DLI and light spectrum (p = 0.001) showed that survival declined more steeply under the shifted spectrum at higher DLI levels. DLI had a significant linear and quadratic effect on CCA growth (p < 0.001), indicating that higher and lower DLI levels inhibit CCA growth. Under the shifted spectrum, turf algal growth was significantly lower with increasing DLI compared to the broad spectrum (p = 0.015). There were no indirect effects of CCA (p = 0.189) or turf algal growth (p = 0.123) on coral survivorship.

**Fig. 3.**
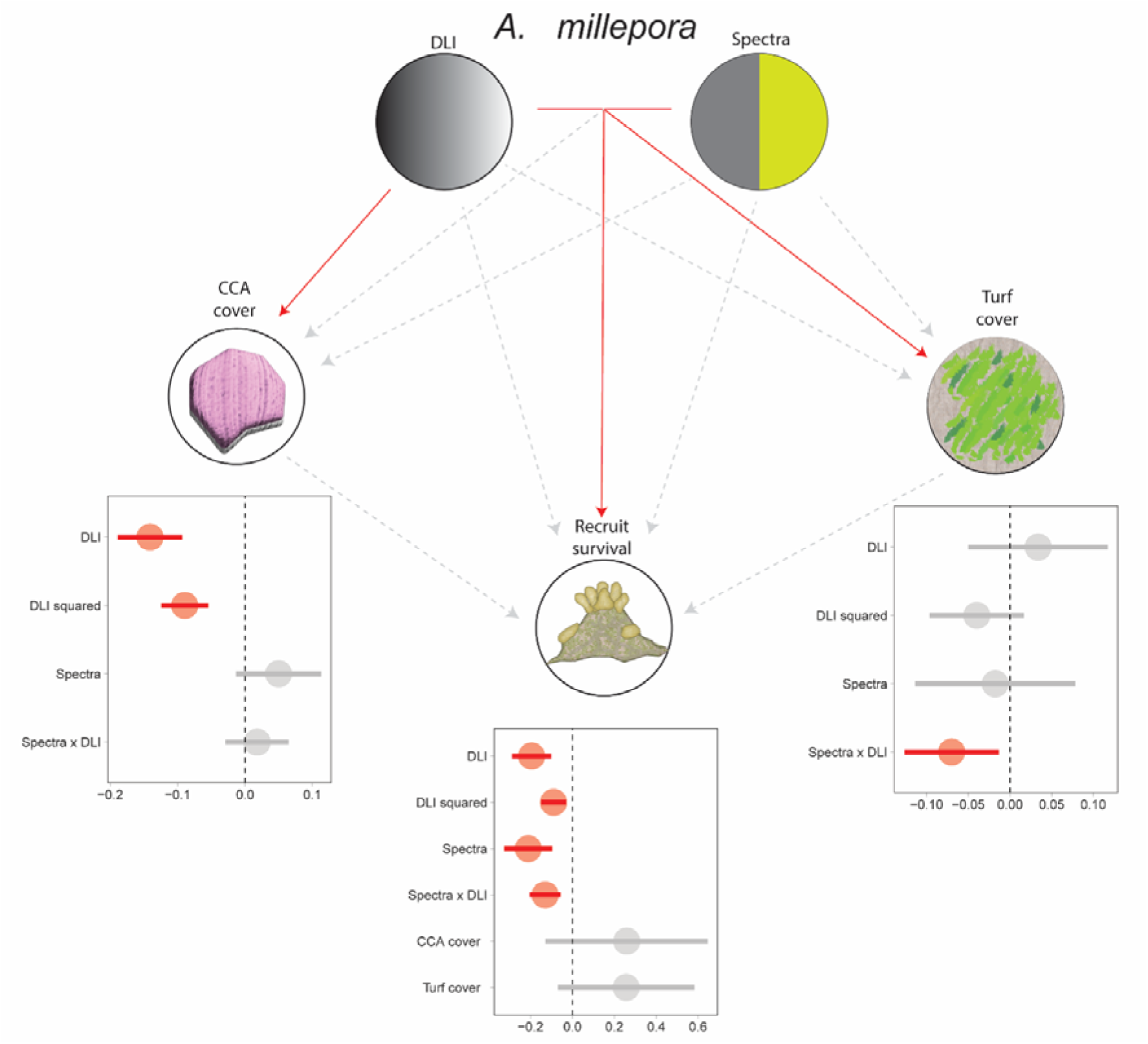
Path diagram of SEM describing the direct and indirect effects of light intensity (DLI), spectra, and the substrate community on *A. millepora* recruit survival. Solid red arrows indicate significant negative pathways, and grey dotted arrows show nonsignificant pathways. Only the interactions are plotted as solid if an interaction occurs. Plots show significant positive (green) and negative (red) scaled estimate and error results for each factor or interaction.

There was a strong inverted U-shaped response of *A.* cf. *tenuis* coral recruit survival under the hormesis model in both the broad-spectral profile (maximum survival = 0.28 mol photons m^-2^ d^-1^, lower EC_10_ = 0.12, upper EC_10_ = 0.61, 95% CI: 0.05–1.62) and shifted-spectra profile (maximum survival = 0.52 mol photons m^-2^ d^-1^, lower EC_10_ = 0.20, upper EC_10_ = 1.02, 95% CI: 0.08–1.80) (Fig. 2c). Maximum growth occurred at the lower light levels (maximum = 1.82 mol photons m^-2^ d^-1^, lower EC_10_ = 0.16, upper EC_10_ >8.72, 95% CI: 0 to >8.72) for the broad-spectral profile with 3-fold growth over the exposure period. However, growth in the shifted-spectral profile remained relatively consistent across all light treatments and had lower levels of growth (Fig. 2d).

The SEM results for *A.* cf. *tenuis* indicated that the model structure aligned well with the observed data (p = 0.554), indicating a good overall fit (Fig. 4). There was no significant interaction between DLI and light spectrum (p = 0.499), or light spectrum only (p = 0.333) on the survival probability of coral recruits. However, there was an effect of both the linear DLI term (p < 0.001) and its quadratic term (p < 0.001) on recruit survival, indicating a significant nonlinear relationship. The cover of CCA was influenced by the quadratic DLI term (p = 0.003), but the effect of light spectrum was not significant (p = 0.715). Turf algal cover showed a positive relationship with the linear and quadratic DLI terms (p < 0.001), but there was a non-significant effect for the light spectrum (p = 0.070). There were no significant indirect effects of CCA (p = 0.749) or turf algal growth (p = 0.252) on coral survivorship.

**Fig. 4.**
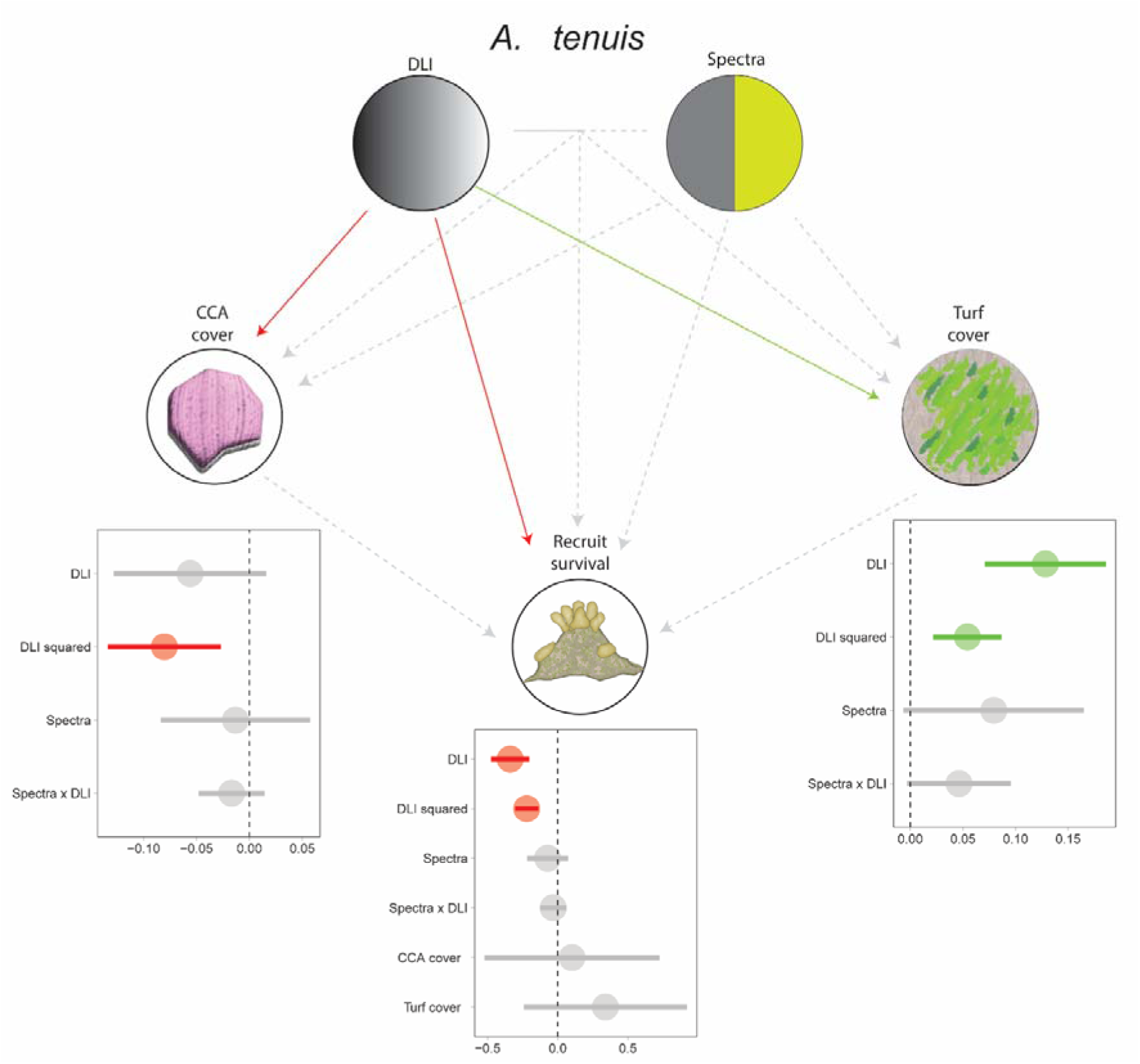
Path diagram of SEM describing the direct and indirect effects of light intensity (DLI), spectra, and the substrate community on *A.* cf. *tenuis* recruit survival. Solid green arrows indicate significant positive pathways, solid red arrows indicate significant negative pathways, and grey dotted arrows show nonsignificant pathways. Only the interactions are plotted as solid if an interaction occurs. Plots show significant positive (green) and negative (red) scaled estimate and error results for each factor or interaction.

The hormesis model showed a marked decline in sponge recruit survival of *P. foliascens* over the 6-week exposure at the higher light intensities, which was more pronounced in the broad-spectral profile. Maximum survival occurred at 0.80 mol photons m^-^² d^-^¹ in the broad-spectral profile (EC_10_ = 2.27, 95% CI: 0–2.86) and at 0.26 mol photons m^-^² d^-^¹ in the shifted-spectral profile (EC_10_ = 1.95, 95% CI: 0–4.05) (Fig. 2e). Growth peaked at low light intensities (broad: 0.34 mol photons m^-2^ d^-1^, shifted: 0.44 mol photons m^-2^ d^-1^ (Fig. 2f). Partial mortality (i.e., alive individuals with partial tissue loss) was observed in some coral and sponge recruits, leading to decreases in relative growth, resulting in values <1.

For the SEM analysis of sponge survivorship and its substrate community, the model structure was consistent with the observed data (p = 0.081) (Fig. 5). Sponge survivorship was significantly related to both the linear (p < 0.001) and quadratic (p < 0.001) terms of DLI, indicating a non-linear relationship. In contrast, light spectrum was not quite significantly associated with sponge survivorship (p = 0.055), and its interaction with DLI was also non-significant (p = 0.128). Similarly, CCA cover showed a significant relationship with both the linear (p <0.001), and quadratic DLI terms (p <0.001), and there was a positive influence by the light spectrum (p = 0.019). For turf algal cover, there was a significant interaction of DLI and light spectrum (p = 0.026). Additionally, there were no indirect effects of CCA growth (p = 0.737) or algal growth on sponge survivorship (p = 0.377).

**Fig. 5.**
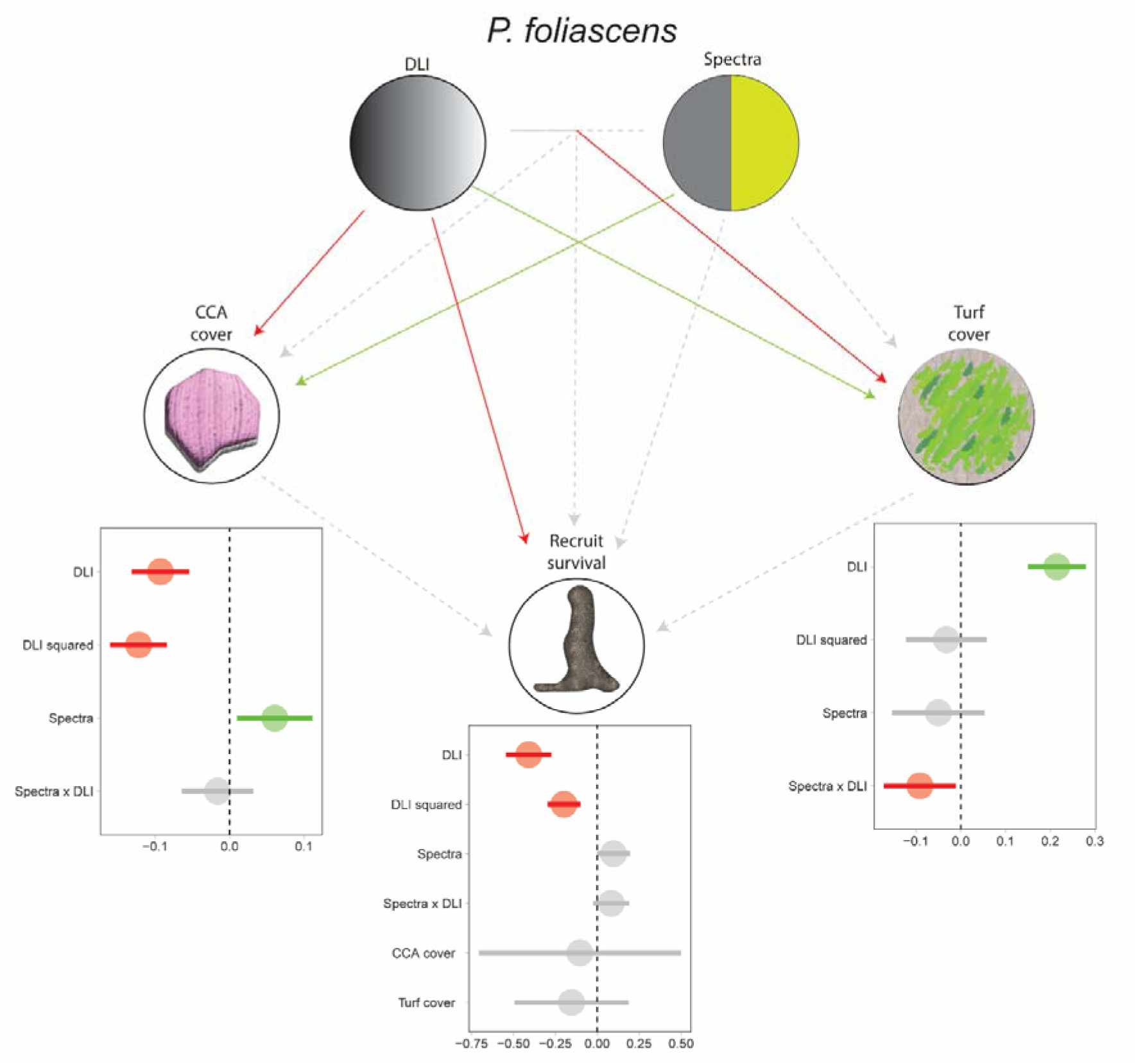
Path diagram of SEM describing the direct and indirect effects of light intensity (DLI), spectra, and the substrate community on *P. foliascens* recruit survival. Solid green arrows indicate significant positive pathways, solid red arrows indicate significant negative pathways, and grey dotted arrows show nonsignificant pathways. Only the interactions are plotted as solid if an interaction occurs. Plots show significant positive (green) and negative (red) scaled estimate and error results for each factor or interaction.

### Impacts on substrate community

For the coral recruit substrates, there was an inverted U-shaped response on CCA cover with light intensity, with the CCA cover peaking (EC_10_: broad = 0.13–0.80 mol photons m^-2^ d^-1^, shifted = 0.11–0.55 mol photons m^-2^ d^-1^) in the low–mid-range light intensities for both spectral profiles (Fig. 6a). Sponge recruits exhibited a similar inverted U-shaped response, with a peak occurring at a slightly higher light intensity (EC_10_: broad = 0.15–0.41 mol photons m^-2^ d^-1^, shifted = 0.31–0.67 mol photons m^-2^ d^-1^) (Fig. 6b).

**Fig. 6.**
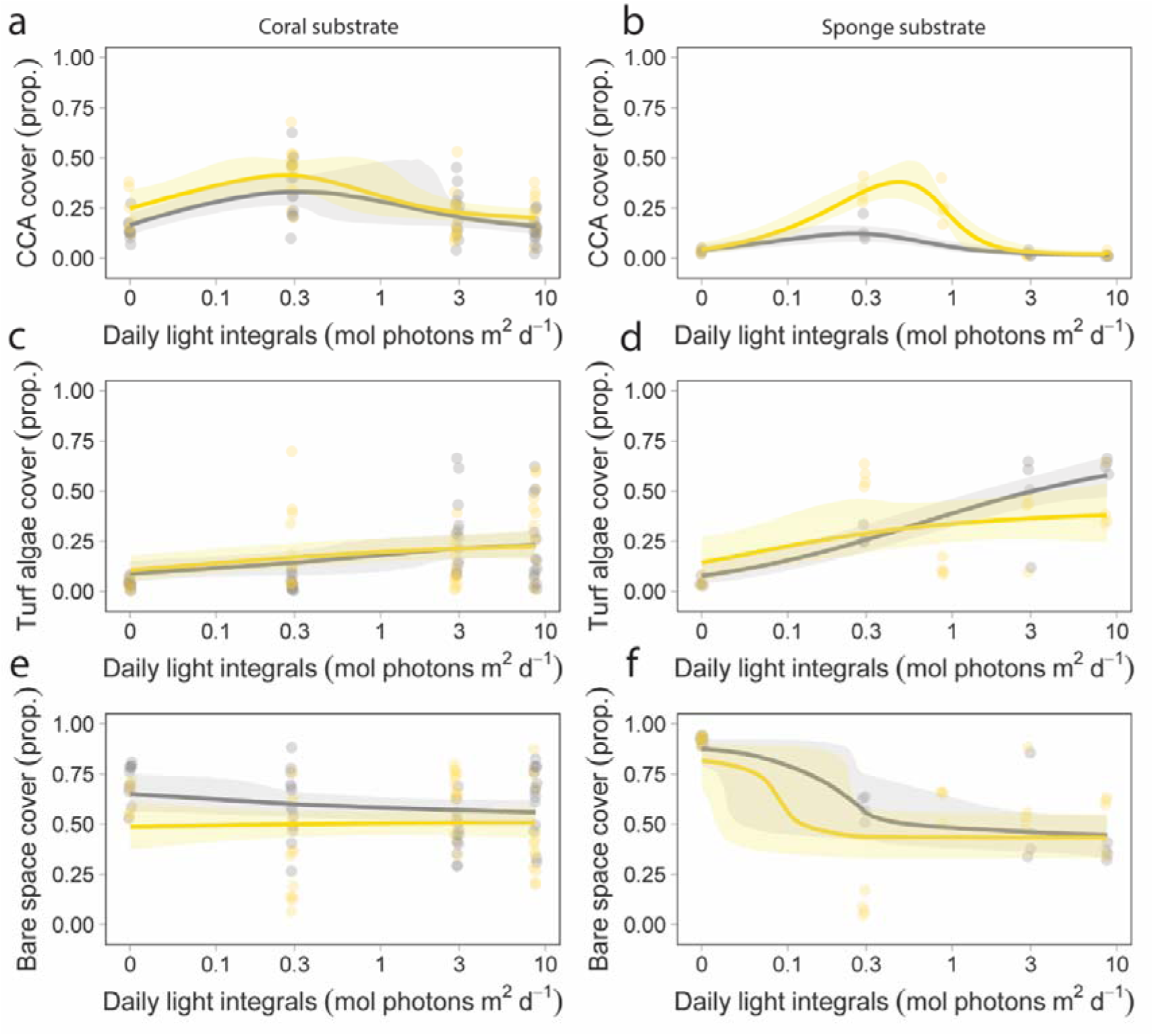
Substrate community composition of the coral-recruit discs (PVC: a,c,e) and sponge-recruit plugs (aragonite: b,d,f) after six-weeks of light exposure under a broad and shifted spectral profile. (a,b) Crustose coralline algal (CCA) cover, (c,d) turf algal cover, (e,f) and bare space. Grey line = broad-spectrum mean line; yellow line = shifted-spectrum mean line.

Turf algal substrate cover increased linearly with light intensity on coral recruit substrates for both spectral profiles and approached ∼23% of the substrate at the highest DLI of ∼9 mol photons m^-2^ d^-1^ (Fig. 6c). Likewise, for sponge recruit substrates, turf algal cover on the substrate generally increased with light intensity, particularly under the broad-spectral profile, with mean cover exceeding 50% of the substrate at the highest DLI of ∼9 mol photons m^-2^ d^-1^. A lower turf algal cover measurement at ∼1 mol photons m^-2^ d^-1^ in the shifted-spectrum treatment was possibly due to recent grazing of snails on some of the substrate.

Bare space, which includes green biofilm-like cover, on the substrates was relatively high, with a small decrease in cover at the broad-spectrum treatment and a relatively constant cover in the shifted-spectrum treatments (Fig. 6e). On the sponge recruit substrates, bare space cover was high in the darkness treatment but decreased to ∼50% of the substrates in the other treatments (Fig. 6f).

### Coral photophysiology

There were no consistent effects of the light treatments on the dark-adapted quantum yield (*F_v_/F_m_*) in either species. A marginal non-significant 16% decrease of *F_v_/F_m_* occurred across the range of DLIs for *A. millepora* under the shifted spectrum. In contrast, a slight increase was observed for *A.* cf. *tenuis* at higher light intensities in both spectra (broad = 11.7%, shifted = 10.9%) (Fig. 7a,b).

**Fig. 7.**
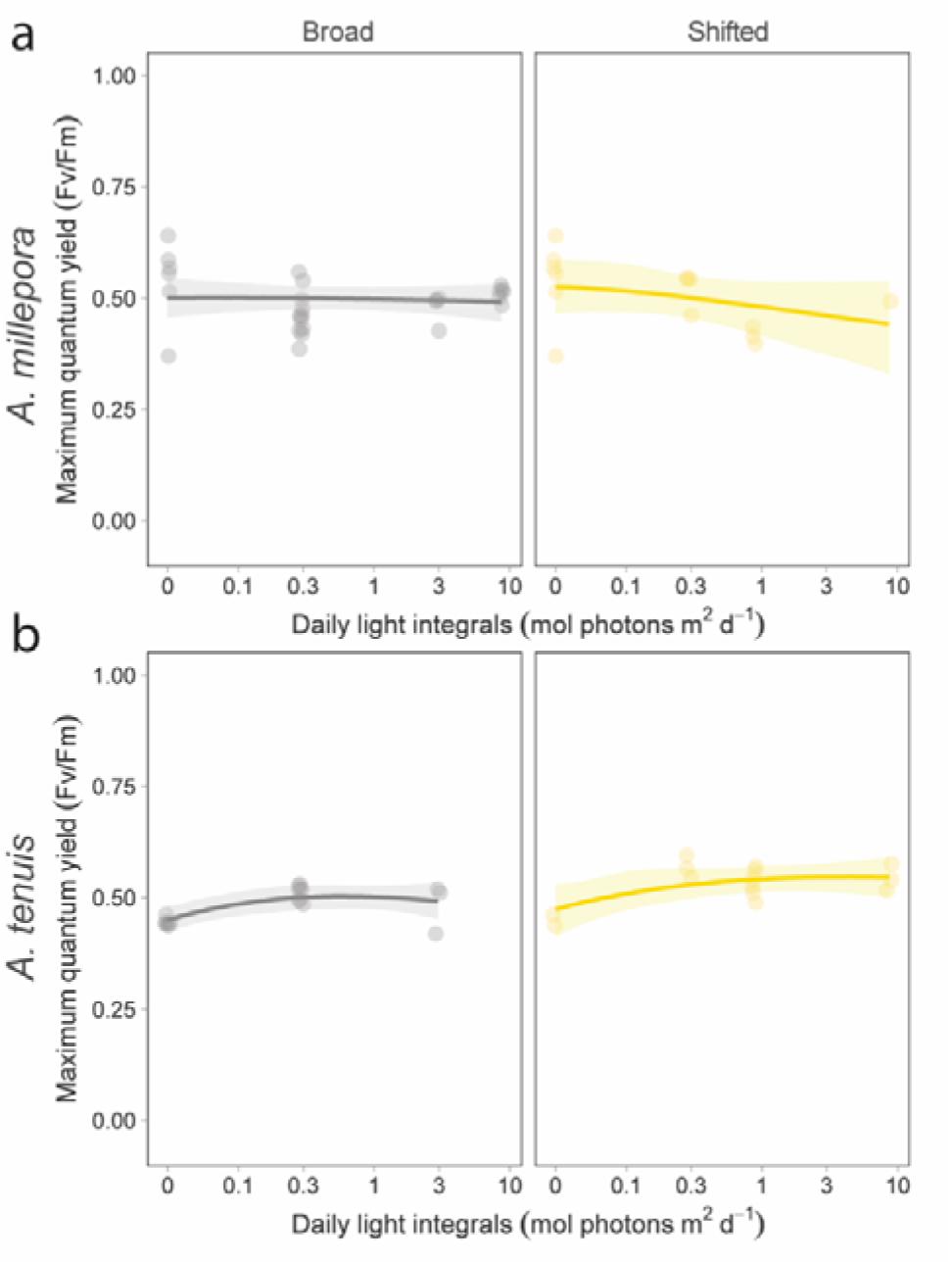
Maximum quantum yields (*F_v_/F_m_*) of coral recruits after the six-week light exposure (broad- and shifted-spectral profiles). (a) *Acropora millepora* and (b) *A.* cf. *tenuis*. Grey line = broad-spectrum mean line; yellow line = shifted-spectrum mean line.

Rapid light curves generate several parameters that indicate the ability of organisms to acclimate to new light conditions (Fig. S2a). *E_k_*indicated that maximum daily experimental irradiance was saturating PSII above ∼50 µmol photons m^-2^ s^-1^ (DLI = ∼2) for both *A. millepora* and *A.* cf. *tenuis* recruits under both spectra (Fig. 8a,c). A decrease in rETR during maximum daily experimental irradiances (signified by *Em*) occurred above ∼200 µmol photons m^-2^ s^-1^ (DLI = ∼4) (Fig. 8b,d). The capacity of *A. millepora* and *A.* cf. *tenuis* recruits to use higher light intensities efficiently reflects the conditions in which they were grown during the first six weeks following settlement, with recruits exposed to high light resulting in increased *rETR_max_* values (Fig. S2b,d). Photosynthetic efficiency (α) appeared to slightly decrease in corals grown in high-light treatments (Fig. S2c,e).

**Fig. 8.**
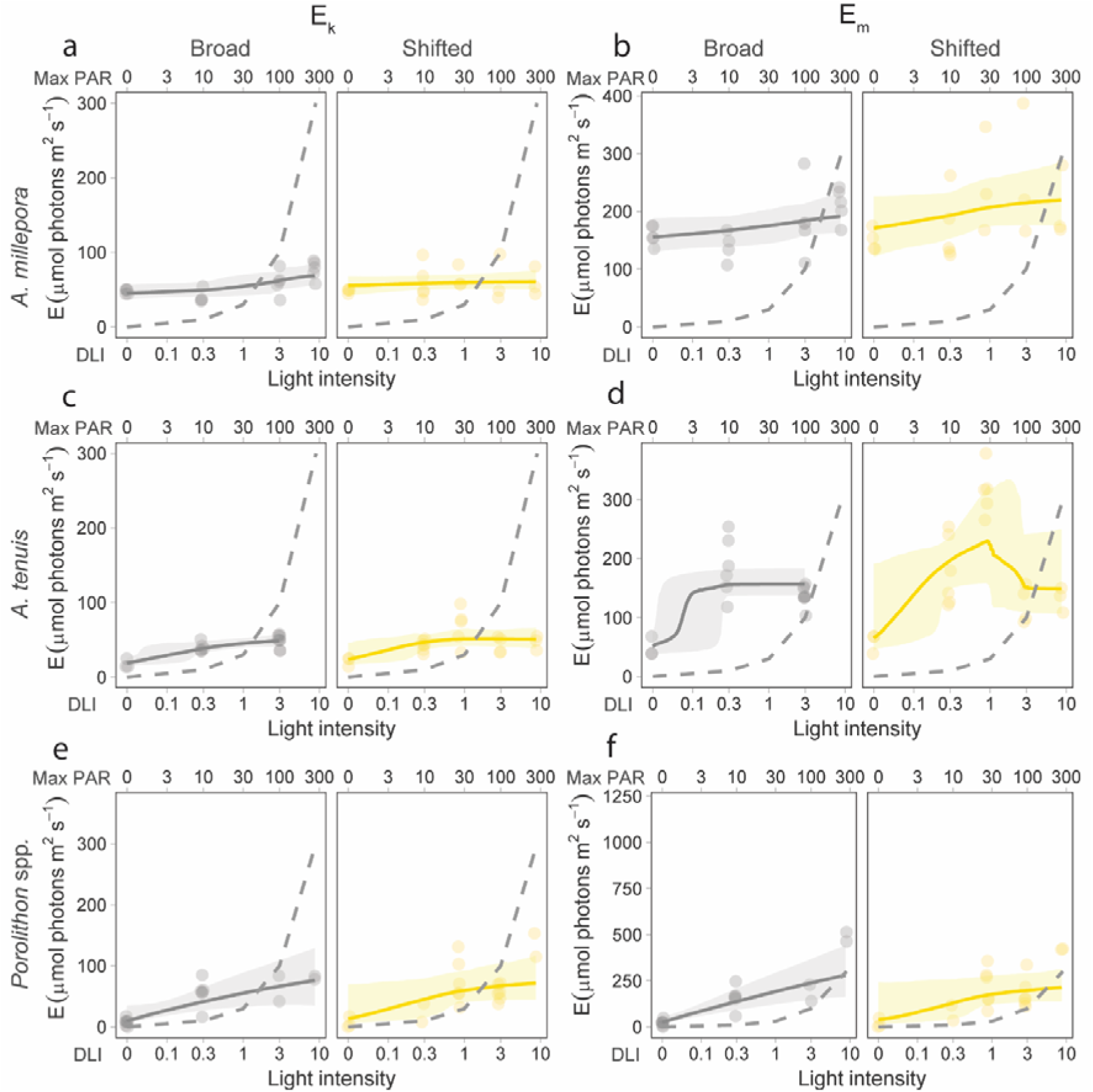
Photobiology of *A. millepora* (a,b), *A.* cf. *tenuis* (c,d), and crustose coralline algae *Porolithon* spp. (e,f) after six-weeks of light exposure under a broad and shifted spectral profile. Rapid light curves parameters *E_k_* (a, c, e), and *E_m_* (b, d, f) following light exposure. *E* denotes irradiance (μmol photons m^-2^ s^-1^). Grey and yellow lines indicate the mean photophysiological responses under broad- and shifted-spectrum light, respectively. Dashed line = maximum daily light intensities during the exposure period.

Under rapid light exposures, all corals for both species utilised similar [1 – Q] and [1 – C] pathways at relatively low actinic light levels (Fig. S5). A consistent trend was that individuals exposed to low-light treatments (<2 DLI) preferentially used photochemical quenching, whereas coral recruits exposed to higher light treatments followed a more typical quenching profile where [1 – Q] and [1 – C] decline with increasing actinic light. The differences between light spectra treatments were relatively similar, although the slightly greater variability in [1 – Q] at higher rapid light exposures in the broad spectrum compared with the shifted spectrum (*A. millepora*: 0.092 and 0.046, respectively; *A.* cf. *tenuis*: 0.120 and 0.065, respectively).

Rapid light curve analysis of the CCA *Porolithon spp.* revealed some acclimation to higher light intensities. (Fig. 8e,f; Fig. S2f,g). *E_k_* increased with light intensity and intercepted with the maximum midday irradiance at ∼75 µmol photons m^-2^ s^-1^ (2–3 DLI), indicating saturation of PSII above these light intensities. The capacity of these CCA to efficiently utilise light was typically better than that of the coral recruits.

## Discussion

Many of the responses to light intensity followed an inverted U-shaped trend, indicating a narrow low-light intensity range that would maximise fitness (i.e., survival, growth, substrate cover). Inverted U-shaped relationships (also referred to as unimodal, *sensu* Gil (2013)) often exhibit an optimum, a relationship commonly observed between organisms and environmental factors. However, the limited number of treatment levels used in many experimental designs often precludes their detection (Gil 2013). Recently, there has been increased interest in identifying environmental optima for corals, known as performance curves, to assess their adaptive capacity across a range of environmental conditions (Jurriaans & Hoogenboom 2019, Silbiger et al. 2019, Álvarez-Noriega et al. 2023), and such curves are also relevant for light characteristics, which likely vary depending on life stage and possibly the light spectrum. Coral and sponge recruit survivorship was influenced by light intensity showing more pronounced negative responses at increased light intensities, and some species- or taxa-specific differences under broad versus shifted spectral profiles. Although these trends occurred across numerous taxa, the inverted U-shaped trend was most pronounced in the survival data, with *A. cf. tenuis* showing the sharpest peak and *P. foliascens* the weakest. Our results suggest that these species may have an optimum low-light threshold above which mortality begins to occur, as reported for other species and life stages (Abrego et al. 2012, McMahon 2018, Kreh 2019, Kuanui et al. 2020).

Only *A.* cf. *tenuis* exhibited strong low-light responses at and near darkness (< 1 DLI) during this early post-settlement stage, while the responses of *A. millepora* and *P. foliascens* were more modest. The lack of substantial light-limitation responses may indicate that recruits primarily rely on residual lipid stores and/or heterotrophic feeding in the first few weeks following settlement to meet their energetic needs (Figueiredo et al. 2012, Conlan et al. 2017). Indeed, recruits can remain in dormancy – a state of low metabolic activity – that may extend for several months under suboptimal environmental conditions (Vermeij & Sandin 2008, Doropoulos et al. 2022). Adult corals and sponges often exhibit strong negative responses to low light (DeSalvo et al. 2012, Pineda et al. 2016, Bessell-Browne et al. 2017, Luter et al. 2021), likely due to their reliance on well-established and stable populations of photoautotrophic microsymbionts, a characteristic not completely developed in these early-stage recruits. The apparent lack of a low-light response in *P. foliascens* recruits may reflect a high degree of photoacclimation by its specialised cyanobacterial symbionts, or an early reliance on heterotrophic nutrition over autotrophy. This could delay sensitivity to light limitation until later developmental stages, once symbiont contributions become more metabolically significant. Recently, Brunner et al. (2022) demonstrated that a light-attenuation period applied to two-month-old coral recruits, rather than those exposed at 1-month of age, caused latent effects on recruit survival, potentially indicating a greater role autotrophy for this later age class. More work is needed to investigate the break-down in symbiosis under extended low-light events, and explore how heterotrophic feeding may mitigate these effects. Regardless, the tolerance of both corals and sponges to low light levels, even under sub-optimal spectra indicate that other sediment-related stressors, such as suspended and depositing sediments, may present a greater risk for these early stage recruits, where low levels for shorter exposure durations have been shown to elicit stronger responses (Abdul Wahab et al. 2019, Tuttle & Donahue 2022).

Both coral species, the sponge *P. foliascens* and to some extent CCA cover showed negative effects of high light. Bleaching of coral recruits (i.e., loss of symbionts) was not observed at higher light levels, likely because the maximum light level used did not exceed the threshold reported to induce bleaching in recruits (∼14 DLI; Abrego et al. (2012)). Similarly, dark-adapted quantum yields (*F_v_/F_m_*) did not indicate any clear reduction in photosynthetic efficiency for low- or high-light adapted recruits. However, rapid light curves at six weeks post-settlement revealed that excess light was largely dissipated via non-photochemical quenching across higher light treatments, with some evidence of PSII down-regulation occurring above 2 DLI. Complete photoinhibition was only observed in a few low-light adapted individuals exposed to high-light levels. Nonetheless, such responses, while primarily protective, may carry energetic costs or signal underlying stress, potentially contributing to the observed mortality in some treatments. The mortality observed in these treatments may also reflect alternative mechanisms such as the direct effects of light on the coral host independent of the symbionts, deleterious effects mediated by the symbionts, or indirect influences on benthic competitors (discussed below) (Huffmyer et al. 2025). The vulnerability of the coral host to high-light intensities may be due to inadequate photoprotection in the form of symbiont xanthophylls, fluorescent proteins (green, cyan, and red fluorescent proteins) and mycosporine-like amino acids, which are present in larvae but may not yet be at adequate quantities post-settlement (Yakovleva & Baird 2005, Roth et al. 2013). In our study, coral recruits grown at darkness were paler near the mouth of the polyps (Fig. S6), which may reflect low retention of symbionts or low chlorophyll content per symbiont (Cumbo et al. 2018). Low quantum yields (*F_v_/F_m_*) in low-light treatments have been observed in other studies (Bessell-Browne et al. 2017, Zaquin et al. 2019, Jones et al. 2021) and may be due to decomposition and unstacking of thylakoid membranes, leading to reduced electron transport (DeSalvo et al. 2012). However, as we did not observe clear *Fv/Fm* responses in our study, these mechanisms are unlikely to have played a role here. Instead, *Fv/Fm* values were overall low regardless of light levels, as typically reported in coral recruits and cell culture studies (Edmunds & Gates 2004, Yuyama et al. 2016, Quigley et al. 2017, Chakravarti & van Oppen 2018, Cumbo et al. 2018), and is discussed further in Text S1. Additionally, the near-Redfield DIN:DIP ratios and more moderate DOC levels suggest that nutrient conditions were unlikely to have promoted strong or consistent nutrient–light interactions or caused significant photophysiological stress, indicating that nutrients could have influenced, but did not govern, the responses to light (Text S2, Table S1).

The CCA community was mostly dominated by *Porolithon spp.*, which notably occurs across a wide range of irradiances *in situ*, but generally increases in growth at lower light levels (midday PAR: 35–60 µmol photons m^-2^ s^-1^, equivalent to 1–1.8 DLIs) (Lewis et al. 2017). We have observed this *ex-situ*, with CCA recruited on bare substrates being increasingly sensitive to irradiances exceeding 200 µmol photons m^-2^ s^-1^ (personal observation, G. Ricardo), potentially indicating increased photosensitivity to the germling stages (Villas-Boas et al. 2023). In our experiment, CCA cover was greatest at approximately 0.3 to 1 DLI consistently displaying an inverted U-shaped trend with light intensity on both substrates. Conversely, turf algal cover generally increased with light intensities, particularly on aragonite substrates. The differences in algal growth observed between the two substrate types may reflect the amount of bare space at the commencement of the experiment, with the mixed algal and biofilm communities being initially less established on the aragonite sponge substrates compared to the PVC coral substrates. Alternatively, the calcium carbonate structure and rugged texture of the aragonite may have facilitated increased algal recruitment in the higher light intensity treatments.

SEM analysis did not reveal any clear significant indirect effects of CCA or turf algae on coral or sponge recruit survival, despite some individuals observed being overgrown by each group. This result suggests that recruits are not overtly competing with crustose coralline or turf algae, in contrast to other studies that have noted competition (Brunner et al. 2022, Noonan et al. 2022). Alternatively, there was not sufficient taxonomic resolution of each group to distinguish drivers of mortality. For example, less conspicuous groups such as cyanobacteria, benthic diatoms, and pathogenic bacteria may have been present alongside the visually dominant green filamentous algae, complicating the interpretation of our observed patterns (Birrell et al. 2008, Arnold et al. 2010, Cárdenas et al. 2016). The inclusion of turf, biofilms and various communities of CCA can complicate interpretations of manipulative laboratory studies such as this study and elsewhere (Brunner et al. 2022, Noonan et al. 2022). For ∼1 mm^2^ sized recruits, even very minor growth of turf and cyanobacteria or other filamentous microorganisms can interact with the edges of the recruit and influence the microscale water chemistry (Kuffner et al. 2006). However, elimination of such taxa is likely unachievable for week- and month-long experiments, nor is it preferable. More sterile and static experiments, while minimising algal growth, will likely introduce experimental artifacts and potentially remove important processes that co-occur during recruitment such as heterotrophic feeding, waste removal, and biomineralization from dissolved nutrients (Gilis et al. 2014, Conlan et al. 2017).

Yellow-green light is considered less useful for driving photosynthesis in algal symbionts (i.e., lower PUR); however, there were no appreciable differences in coral and sponge responses to these disparate light profiles across most realistic light treatment combinations. Although monochromatic light experiments can provide insights into which wavelengths are contributing survivorship, growth and reproduction in marine taxa (Wijgerde et al. 2014, Strader et al. 2015, Strydom et al. 2017a), the same effects are not necessarily observed under more realistic spectral profiles that include a mix of wavelengths (Strydom et al. 2017b, Jones et al. 2021, Ricardo et al. 2021). During a preliminary study on adult corals, Jones et al. (2021) used a similar shifted profile to that employed here. The authors did not observe any effects on survivorship, and only limited negative effects on growth under a more extreme light intensity and spectral treatment combination (6 DLI at a ∼0.2 blue:green ratio). However, sublethal effects were observed on algal density, Chl *a* content, and lipids. At more environmentally relevant shifted light treatment combinations of 1 DLI at ∼0.2 blue:green ratio, there were little differences observed between the shifted and broad spectrum, indicating responses observed were mostly driven by low light rather than changes in the spectral profile, as we similarly observed. Complementary chromatic adaptation has been shown in cyanobacteria, and it is possible that phycocyanin and phycoerythrin pigments in the cyanobacterial symbionts within sponges can extend the PUR range further towards the centre of the spectrum (∼450–550 nm) (Luter et al. 2021). Subtle spectral effects may also be masked by increased heterotrophic energy acquisition, as observed in corals in turbid environments (Anthony & Fabricius 2000). Further, the actinic light supplied during rapid light curve analyses rarely matches the spectra profiles that marine organisms are acclimated (Schreiber et al. 2012), as was the case in this study; yet the consistency in photophysiological responses between the broad-spectrum and shifted-spectrum treatments suggests that any potential bias from the spectral mismatch likely did not affect the validity of the observed trends. While high light levels in the shifted spectral profiles may have led to increased mortality in *A. millepora* relative to the broad spectra, turbidity shifted-spectral profiles negatively correlate with light intensity, and therefore higher light intensities under shifted-profiles are limited in their environmental relevance and should be interpreted with caution. Even under doldrum conditions and cloud-less skies, blue:green profiles <0.2 are unlikely to correspond to light intensities greater than ∼4 DLI (Fig. 1a,b), and even less so in shaded areas typical of recruit habitat.

Several studies suggest coral larvae settle predominantly in small crevices, similar in scale to their body length (Nozawa et al. 2011, Whalan et al. 2015, Ricardo et al. 2017, Randall et al. 2021). The light conditions within these <2 mm crevices of this scale are unknown because probes of this size are not readily available; however, measurements taken *in situ* suggest that larger, more exposed crevices (∼50 × 50 mm) on vertical surfaces and undersides are regularly subject to low light of <50 µmol photons m^-2^ s^-1^ (Box & Mumby 2007, Briggs 2016, Doropoulos et al. 2016), and for inshore reefs very low light of <10 µmol photons m^-2^ s^-1^ (Ricardo et al. 2021). This indicates, along with the results presented here, that these low-light habitats may initially be optimal for recruit survival and growth. Eventually as recruits grow out from these crevices and undersides and their symbiont community becomes more established, they are exposed and better utilise the exponentially increasing light exposure. However, this trajectory may not hold in persistently turbid environments where sedimentation or coastal pollution limits light penetration, potentially constraining access to light even as corals grow.

During restoration practices, a significant impediment to successful outplanting of corals and other taxa is navigating their recruits through the post-settlement stage (Edmunds et al. 2024), a task complicated by the multiple environmental stressors and changes in physiology that occurs within the first months following settlement. Understanding the micro-environments that these recruits inhabit, alongside interactions with the substrate community is a crucial step in improving recruit grow-out, particularly for inshore reefs where numerous water quality thresholds cooccur (Doropoulos et al. 2016, Randall et al. 2020, Brunner et al. 2021). Here, we show that coral recruits and sponges may benefit from an initial low-light environment that will i) increase survivorship, ii) increase CCA growth and iii) decrease turf algae growth. However, these conditions will not to be beneficial in the mid-to-long-term as mature adults increasingly rely on autotrophy to meet energetic needs (Muscatine 1990, Brunner et al. 2022), and an incremental increase in total light intensity (i.e., DLIs) may provide optimal conditions through the early-life stages. One clearly unresolved question pertains to the specific post-settlement age autotrophy becomes the dominant energetic pathway in corals and phototrophic sponges (Cheshire & Wilkinson 1991, Hazraty[Kari et al. 2022). Furthermore, the possibility of manipulating this process, such as through modifications in the timing, concentration, and diversity of symbionts during the inoculation of recruits, paired with variations in light intensity, to optimise growth and survival, is an avenue yet to be thoroughly explored.

Our results reveal complex relationships between the substrate community and early-stage recruits of both corals and sponges, highlighting distinct low light-intensity optima for each taxon, but less consistent effects of the shifted yellow–green spectral profile. As recruits begin to develop into larger, fully autotrophic juveniles with increasing metabolic demands, their light requirements are likely to rise, which may also increase their vulnerability to turbidity-driven light attenuation associated with coastal darkening. Nevertheless, early-stage recruits appear relatively resilient to extended low-light exposure. Therefore, other water quality stressors with lower known tolerance thresholds, such as suspended and deposited sediments, may pose a greater risk during this early stage and represent a more effective focus for targeted management.

## Acknowledgements

This project was supported through government appropriation funding (AIMS) and from the Australian Government’s National Environmental Science Program (NESP). We acknowledge In-situ Marine Optics for their technical expertise with the light sensors. We acknowledge Paul Boyd and Andrea Severati for design of the lighting systems, and the AIMS SeaSim team for assistance for providing conditioning tanks, room set-up and animal husbandry requirements. We would also like to thank Manuel Maldonado, Florita Flores, Lonidas Koukoumaftsis, Christian O’Dea, and Samantha Crisp for assistance with coral spawning, and David Suggett and Carlos Alvarez Roa for advice on the photophysiology. The authors also acknowledge the Manbarra, Bindal and Wulgurukaba people as the Traditional Owners of the sea country where this work took place.

