## Supplementary Material for "Unravelling the influence of light on inshore coral and sponge recruits and their substrate communities"

Text S1. Infection of early recruits with cultured symbionts is a common method used for *ex situ* coral rearing (Pollock et al. 2017); however, these symbionts maintain lower maximum quantum yields (0.4–0.5 *F_v_/F_m_*) in culture relative to adults corals at (DiPerna et al. 2018), and ~1 yr. old juveniles (Kuanui et al. 2020). While these lower yield values are interesting, and have been observed frequently for coral recruits and symbiont cells in culture (Edmunds & Gates 2004, Yuyama et al. 2016, Quigley et al. 2017, Chakravarti & van Oppen 2018, Cumbo et al. 2018), it is difficult to speculate the mechanism, which is outside the scope of this study. However, it has been found that optically dense, heterogeneous, or thick samples can influence PAM measurements and therefore caution is needed in comparing optically thin samples, such as cell cultures and coral recruits, with adult corals (Wangpraseurt et al. 2019, Serôdio & Campbell 2021). Regardless, early coral recruits hosting cultured symbionts grow at a comparable rate to recruits that acquire symbionts from non-cultured means (Fig. S7), and the *F_v_/F_m_* values were within the range of which typically occurs from symbionts collected from sediment samples across the GBR inshore-offshore reef gradient (Quigley et al. 2017).

Text S2. Dissolved inorganic nitrogen (DIN) and dissolved inorganic phosphorus (DIP) influence both autotrophic and heterotrophic functioning in corals and sponges. DIN and DIP values were overall elevated but remained lower than that reported during most wet season flood plumes. There was no major deviation of mean DIN:DIP ratios (14.0:1 ± 3.8:1) from the Redfield ratio (16:1) (Redfield 1958), therefore neither nitrogen nor phosphorus limitation is likely, or at least, deviations were only temporary. Absolute concentrations of DIN were moderate, particularly following rainfall events, but there was no clear evidence of nutrient-induced photophysiological stress interacting with light: maximum quantum yield (*Fv/Fm*) remained stable under high light treatments, suggesting no damage to PSII. Although elevated DIN has been associated with a range of physiological responses under high light in other studies (Wiedenmann et al. 2013, Rosset et al. 2017), such mechanisms appear unlikely to explain the reduced growth and survivorship observed here.

Additionally, ambient levels of dissolved organic matter (DOM), as indicated by measured DOC concentrations (1.07 ± 0.26 mg L^-1^; Table S1), may have influenced microbial activity or heterotrophic feeding by coral recruits, particularly under low light where phototrophic carbon supply is reduced. However, given the moderate nutrient concentrations recorded, it is likely that light remained the primary driver, with nutrients acting as a potential modulator of the growth/survival responses

Table S1. Water quality of incoming general purpose filtered seawater to the experimental room (mean ± standard deviation)

| **Salinity (ppt)** | **pH** | **Alkalinity (µmol kg⁻¹)** | **DIC (µmol kg⁻¹)** | **DOC (mg L⁻¹)** | **NH₄ (µmol L⁻¹)** | **NO₂ (µmol L⁻¹)** | **NO₃ (µmol L⁻¹)** | **PO₄ (µmol L⁻¹)** | **DIN:DIP ratio** | **SiO₄ (µmol L⁻¹)** |
| --- | --- | --- | --- | --- | --- | --- | --- | --- | --- | --- |
| 35.1 ± 0.7 | 8.12 ± 0.04 | 2318 ± 59 | 2063 ± 57 | 1.07 ± 0.26 | 0.75 ± 1.48 | 0.20 ± 0.11 | 2.49 ± 1.36 | 0.23 ± 0.12 | 14.02 ± 3.77 | 12.3 ± 11.2 |

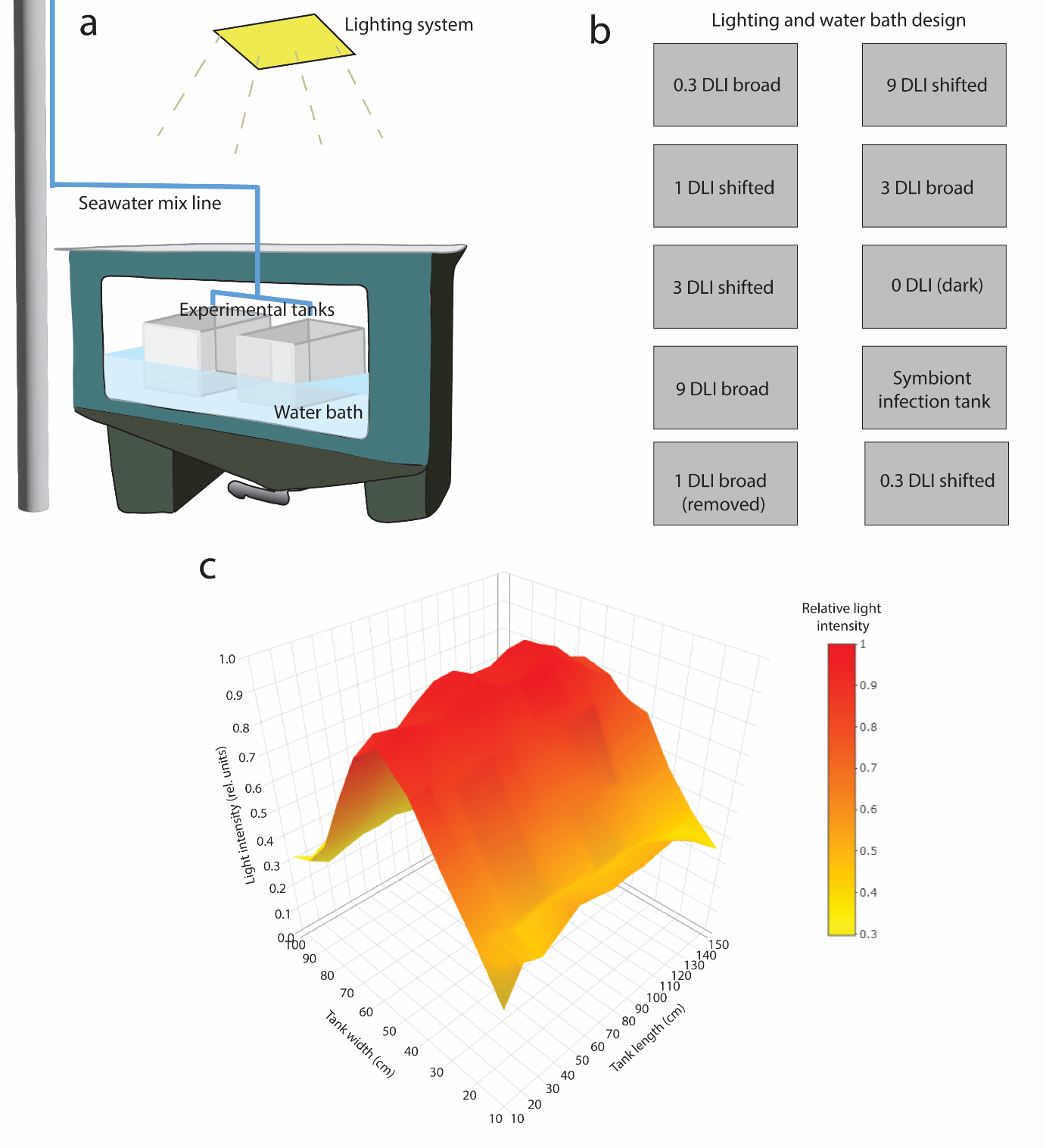

Fig. S1. Experimental design and light intensity distribution. (a) Individual light treatment and water bath set-up. Experimental tanks contained both coral and sponge substrates. (b) Randomised arrangement of light treatments within the experimental room. (c) Light intensity distribution under each lighting system at the level of the 150 × 100 cm water-bath tank.

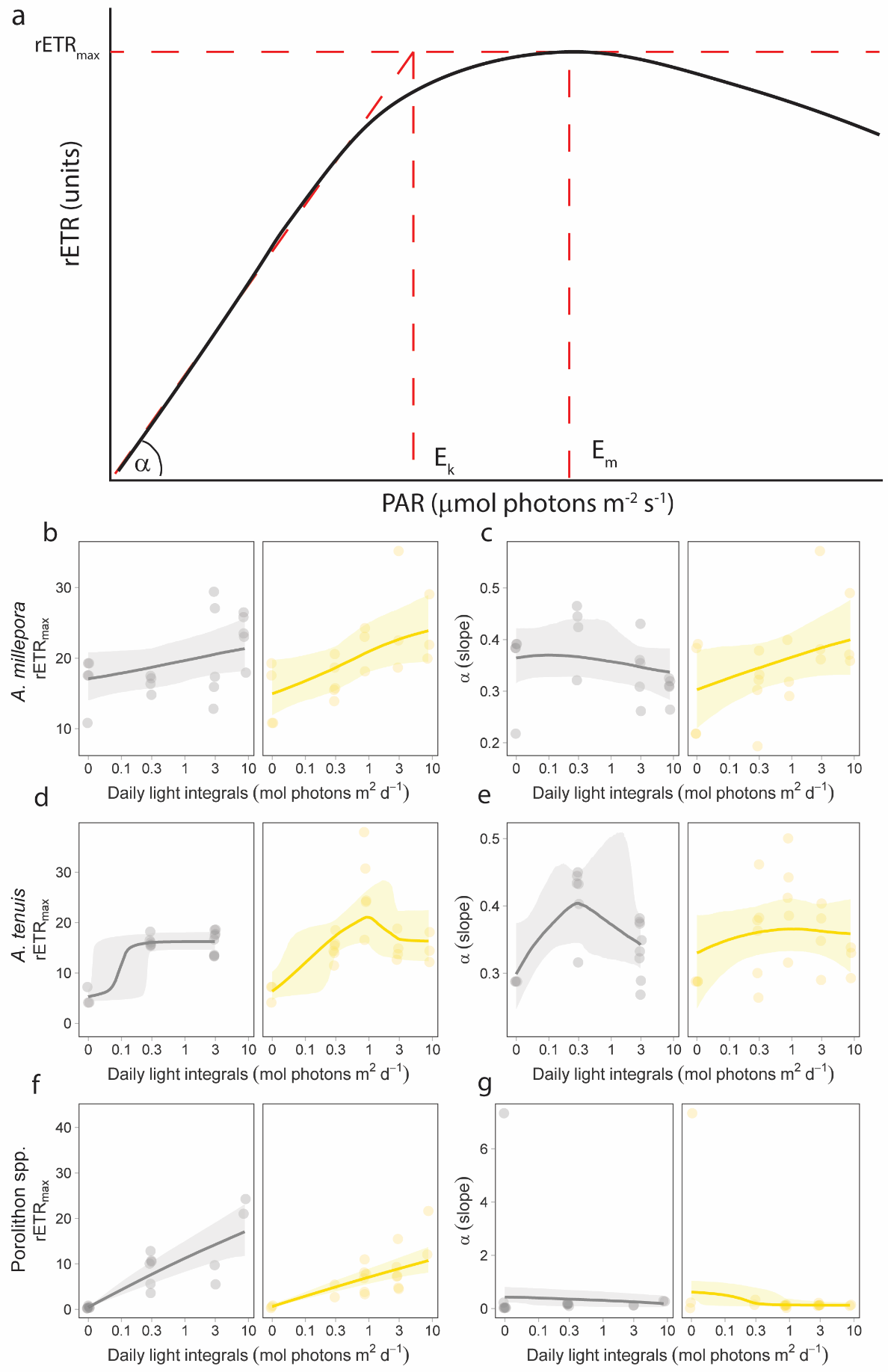

Fig. S2. a) Schematic diagram of a rapid light curve analysis (modified from Ralph and Gademann, 2005). Data are fit with a double exponential decay function (Platt equation) to determine parameters α, *rETR_max_*, *E_k_*, and *E_m_*. *rETR_max_* (b, d, f) and α (slope) (c, e, g) for *A. millepora* (b, c), *A.* cf. *tenuis* (d, e), and *Porolithon* *spp*. (f, g). Grey and yellow lines indicate the mean photophysiological responses under broad- and shifted-spectrum light, respectively

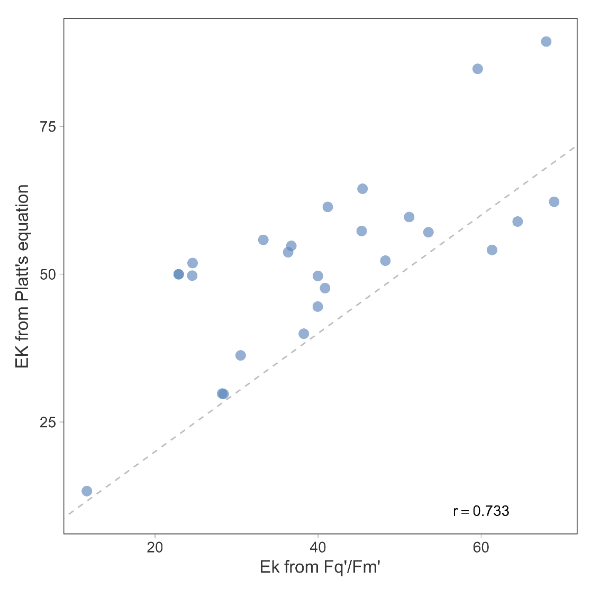

Fig. S3. Correlation between *Ek* derived from Platt equation and *Ek* derived from the light-dependent quantum efficiency of PSII.

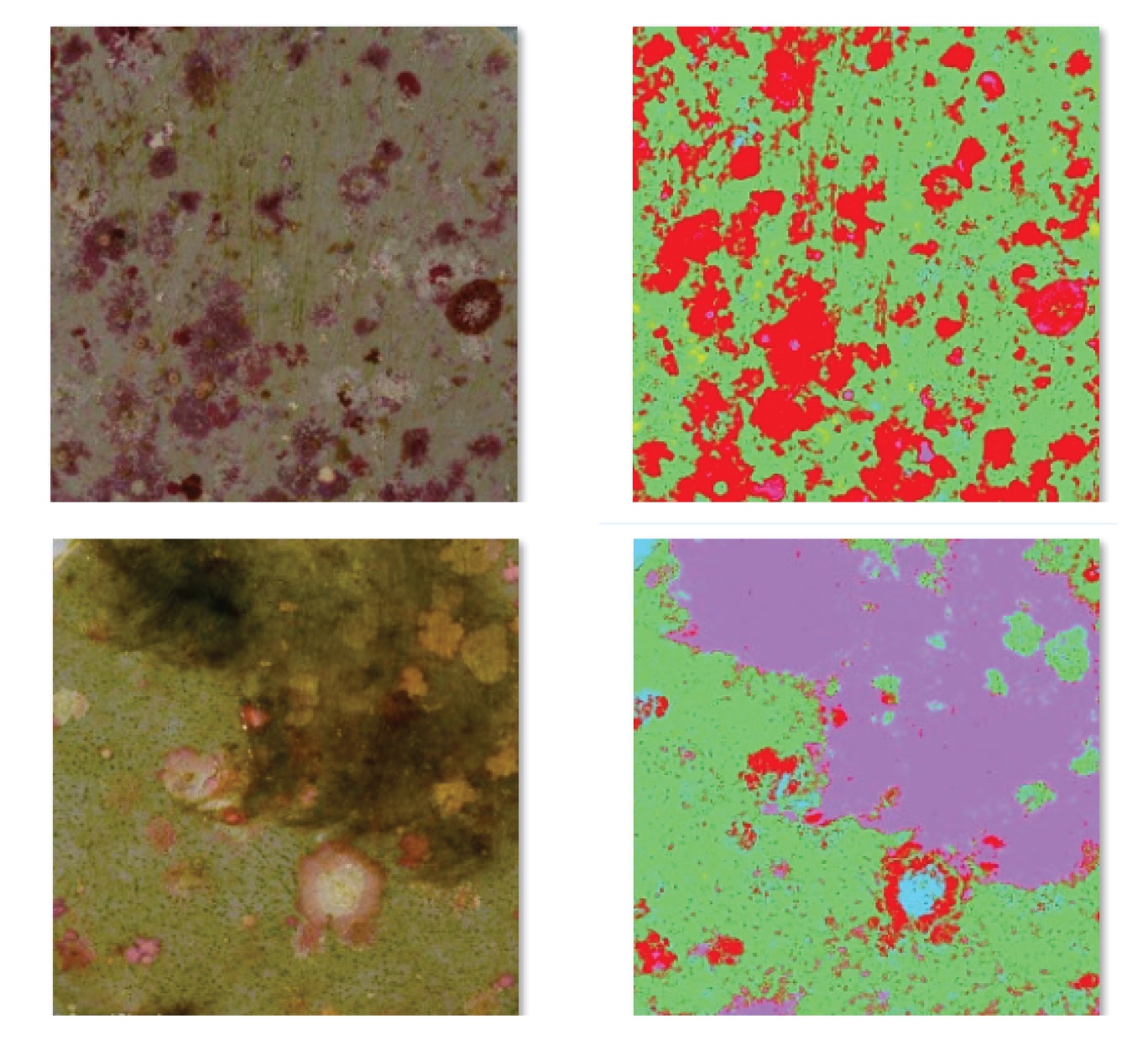

Fig. S4. Two representative images of the segmented coral recruit substrates (PVC). Red = CCA, Green = biofilm. Purple = turf algae. Yellow = bare space. Blue = bleach/glare (removed). Classification was performed using the ImageJ plugin Trainable Weka Segmentation.

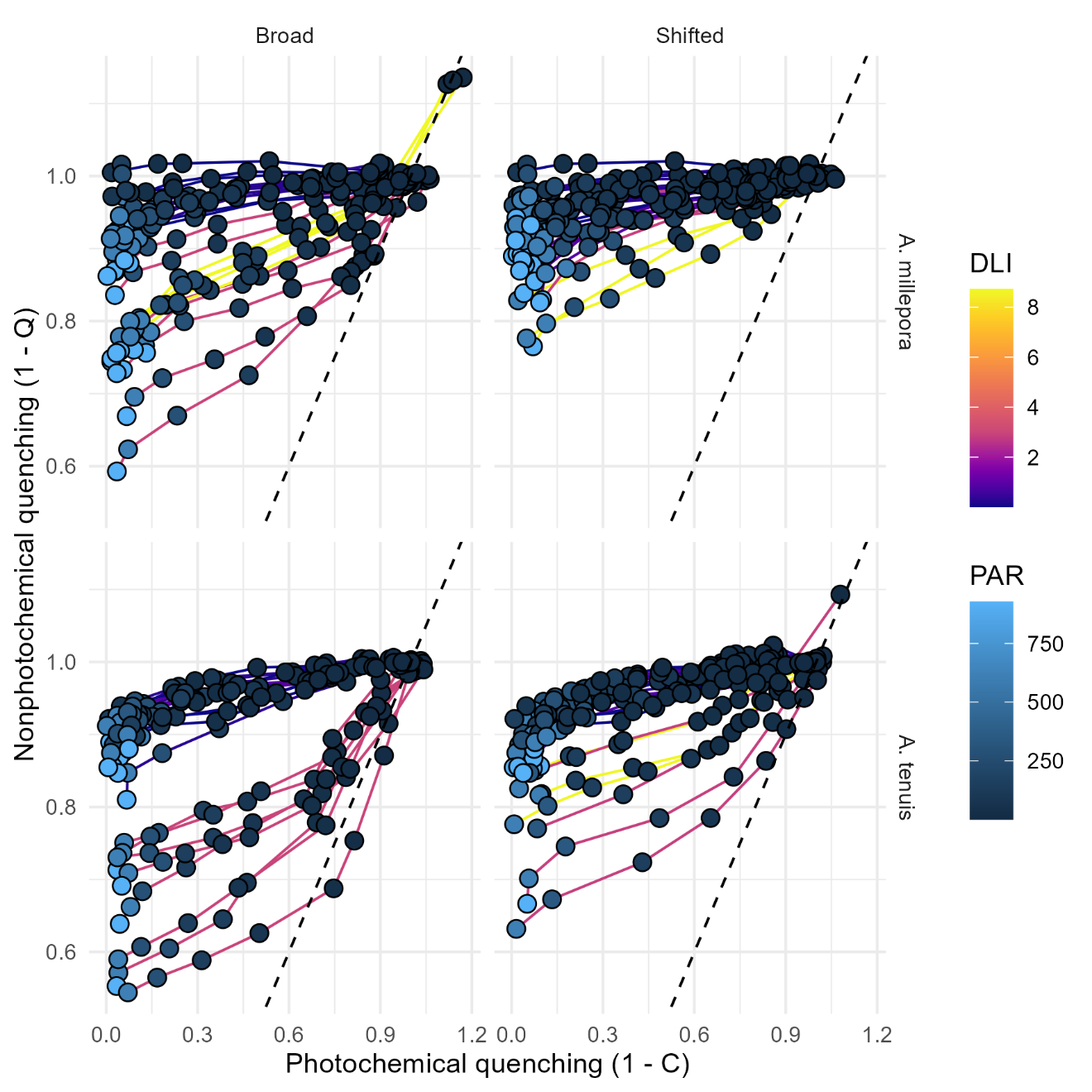

Fig. S5. The relationship between photochemical quenching (1 – C) and nonphotochemical quenching (1 – Q) under broad and shifted spectra for *Acropora millepora* and *Acropora* cf. *tenuis*. Dashed line indicates an equilibrium between each pathway. Connecting lines denote individuals, coloured at their light intensity treatment exposure. Coloured points denote the actinic light of the rapid light curves.

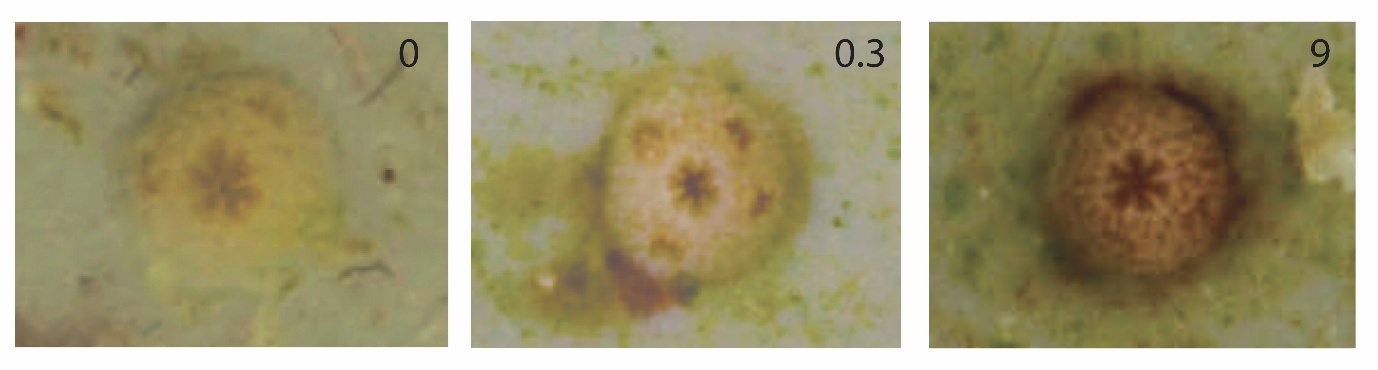

Fig. S6. Representative images of coral recruits exposed to broad-spectrum light at 0, 0.3, and 9 daily light integrals (DLI). Tissue pigmentation serves as a visual indicator of symbiont status, with darker recruits reflecting successful symbiont uptake and retention, and paler individuals indicating reduced uptake or partial loss of symbionts.

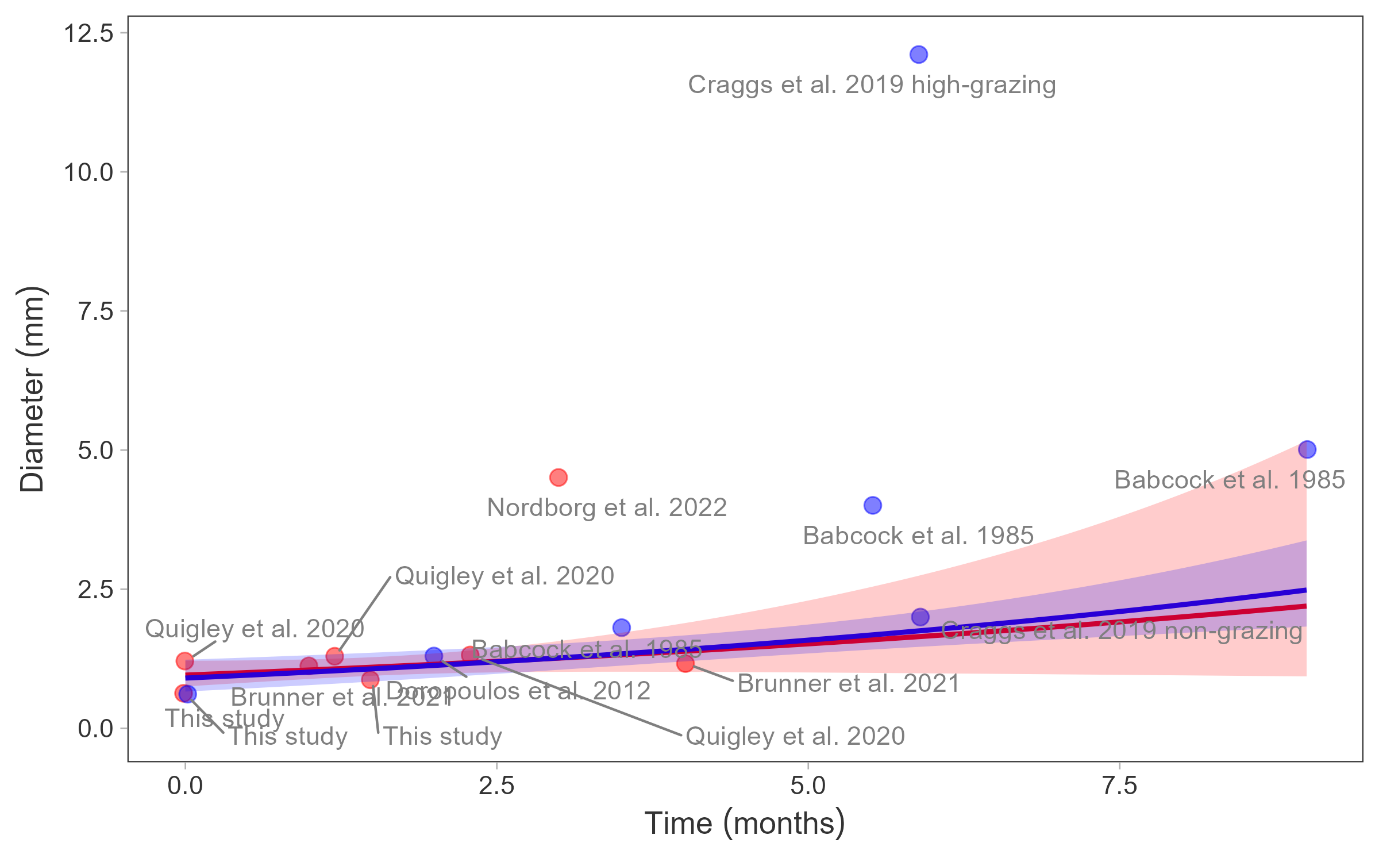

Fig. S7. Coral recruit growth rates from several studies including this study using symbiont infection of cultured symbionts (red), and symbionts infections acquired from non-cultured symbionts (blue). Growth data from this extracted from Babcock (1985), Doropoulos et al. (2012), Craggs et al. (2019), Quigley et al. (2020), Brunner et al. (2021), Nordborg et al. (2022). A log-normal generalised linear model was fit to the data. Shading represents the 95% CI.
